# Progressive Alignment of Inhibitory and Excitatory Delay May Drive a Rapid Developmental Switch in Cortical Network Dynamics

**DOI:** 10.1101/296673

**Authors:** Alberto Romagnoni, Matthew T. Colonnese, Jonathan D. Touboul, Boris Gutkin

## Abstract

Nervous system maturation occurs on multiple levels, synaptic, circuit, and network, at divergent time scales. For example, many synaptic properties mature gradually, while emergent network dynamics, as data show, change abruptly. Here, we combine experimental and theoretical approaches to investigate a sudden transition in spontaneous thalamocortical activity necessary for the development of vision. Inspired by *in vivo* measurements of time-scales and amplitudes of synaptic currents, we extend the Wilson and Cowan model to take into account the relative onset timing and amplitudes of inhibitory and excitatory neural population responses. We study the dynamics of this system and identify the bifurcations as the onset timescales of excitation and inhibition are varied. We focus on the specific typical developmental changes in synaptic timescales consistent with the experimental observations. These findings argue that the inhibitory timing is a critical determinant of thalamocortical activity maturation; a gradual decay of the *ratio* of inhibitory to excitatory onset time below one drives the system through a bifurcation that leads to a sudden switch of the network spontaneous activity from high-amplitude oscillations to a non-oscillatory active state. This switch also drives a marked change to a linear network response to transient stimuli, agreeing to the *in vivo* observations. The switch observed in the model is representative of the sudden transition in the sensory cortical activity seen early in development.

## Introduction

During development, complex biological processes combine to generate, shape and refine the nervous system. In the developing mammalian brain, particularly notable changes in thalamocortical activity patterns occur eventually leading to a functional circuit. These intermediate thalamocortical activity regimes reportedly reflect the progression of the circuits through configurations specialized for developmental roles enhancing synaptic plasticity, amplification, and conservation of energy, and precede a sudden transition to a functional network (Luhmann et al. 2016; Kanold and Luhmann 2010; Kirkby et al. 2013; Butts and Kanold 2010; Khazipov et al. 2013).

While the evolution of a variety of properties from the cellular to the synaptic and circuit scales throughout development have been well described, how these maturations lead to changes in the emergent collective activity patterns of brain networks remains to be established. Of particular interest is an apparent complex and, at times, unclear relationship between the microscopic circuit properties, which tend to gradually progress from immature to adult levels (McCormick and Prince 1987; Luhmann and Prince 1991; Etherington and Williams 2011), and the macroscopic patterns of activity, whose transitions can be abrupt (Colonnese et al. 2010; Rochefort et al. 2009; Golshani et al. 2009; Chipaux et al. 2013). An important question in development is whether such sudden “switches” in network dynamics reflect dramatic changes in the electrophysiological properties of cells or network circuits, or arise as nonlinear responses of the macroscopic activity to gradual changes in microscopic parameters. Establishing the relationship between cellular/synaptic maturation and the emergence of new activity patterns is thus a critical component for understanding how neural circuitry becomes functional. It is also important in order to predict how neurological disorders, which often cause subtle cellular and synaptic changes, might have outsize effects during the important developmental epochs when synapses and circuits are forming (Ackman and Crair 2014; Ben-Ari 2008).

Delineating how slow developmental changes at the microscopic scale relate to the fast, macroscopic transitions is a challenge for experimental approaches. Inhibiting even processes deemed non-essential can have significant effects on the network dynamics, making it particularly challenging to disentangle the respective roles of the multiple synaptic and cellular changes in the rapid functional transitions. This is where modelling and mathematical analysis becomes essential. Sudden changes in the activity dynamics caused by smooth changes in the underlying parameters are common properties of recurrent networks (and of non-linear dynamical systems in general). Mathematically, these can be identified through bifurcation theory, that classifies changes in the number or stability of attractors occurring in dynamical systems as parameters are varied. Applications of bifurcation theory to neurosciences are abundant: they were used to account for the emergence of visual hallucinations due to cortical disinhibition (Ermentrout and Cowan 1979), perceptual bistability (Shpiro et al. 2009), neuronal bursting caused by slow adaptation of cellular excitability (Ermentrout and Kopell 1986; Izhikevich 2000), and epileptic-like activity in response to slowly increasing input levels (Touboul et al. 2010).

In this paper we apply an analytic approach to identify bifurcations in a reduced neural population activity model to show that a gradual maturation of inhibitory time-scales can account for a dramatic developmental change in cortical activity: the sudden emergence of an asynchronous thalamocortical activity regime that allow for linear responses to sensory input (Colonnese and Phillips 2018). In rat visual cortex, this switch in thalamocortical network dynamics affects both spontaneous as well as light-evoked activity. It consists of a massive reduction in total excitability of the network as the systems shifts from a developmental-plasticity mode, into a linear sensory coding mode correlated with the emergence of the ability for the animal to cortically process visual stimuli, all in under 24 hours. Day-by-day measurements in somatosensory and visual cortices have revealed a number of synaptic and circuit changes that might contribute to this switch in cortical activity. These include an increase in feedforward inhibition triggered in both cortex and thalamus resulting in reduced inhibitory delay (Colonnese 2014; Murata and Colonnese 2016; Minlebaev et al. 2011; Daw et al. 2007), as well as more gradual increases in total inhibitory and excitatory synaptic amplitudes, increasing inhibitory effect due to reduction in the reversal potential for chloride (Owens et al. 1996; Glykys et al. 2009), changes in interneuron circuitry (Tuncdemir et al. 2016; Marques-Smith et al. 2016), patterning of thalamic firing (Murata and Colonnese 2016; Lo et al. 2002; McCormick et al. 1995; Murata and Colonnese 2018), reduction of action potential threshold (Golshani et al. 2009), and changing glutamatergic receptor composition (Rumpel et al. 2004), among others.

Since inhibitory interneurons play a critical role in determining the network properties of adult thalamocortical circuits, particularly the oscillatory synchronization and bi-stability (Haider and McCormick 2009), we hypothesized that changes in inhibition, particularly in the timing of inhibition, is a likely determinant of the rapid emergence of functional visual circuits. While previous modelling approaches studied the role of inhibitory offset timing (Ermentrout 1998; Destexhe and Sejnowski 2009), their use of order-one differential equation models precluded analysis of the onset timescales which are most relevant to development. To enable us to analyse the role of changing inhibitory/excitatory strength and onset delay (two critical parameters whose impact was largely overlooked in the computational literature) on network activity, we developed a novel extension of the Wilson and Cowan (WC) model (Wilson and Cowan 1972), in which the ratio of excitatory to inhibitory current amplitudes, as well as relative timing can be varied independently to determine their combined potential roles in the evolution of cortical activity.

In this work, we first review the key experimental observations that serve as a foundation of our computational study, and that our model must capture. We then introduce our extension of the WC firing-rate model, and identify key parameters that will serve in the analysis, notably, the ratio in the excitatory-to-inhibitory onset time-scales, the ratio of the excitation-to-inhibition strength and the amplitude of the external stimulus. This model being developed, we then carry out an extensive codimension-one and -two bifurcation analysis of the model to identify the critical changes in the dynamics of the system determined by the key parameters of input strength and relative inhibitory amplitude and delay. By using these results we are able to identify developmental parameter trajectories that are consistent with the experimental data during the transition in network activity.

Our results lend support to the dynamical hypothesis that dramatic changes in the activity of neural networks during development do not require similarly sharp changes in the cells’ or network’s properties. It also suggests that, for the case of the rat visual cortex, a speed-up in the time-scale of the inhibitory neuronal response onset can account for multiple key observations on cortical activity throughout development, including the sudden switch from oscillatory to an active stable network behavior similar to that observed in vivo as well as a switch from all-or-none bursts evoked by large transient inputs, to a linear response to stimuli reflecting their amplitude and duration.

## Material and Methods

In all species and systems examined, sensory thalamocortical development can be broadly divided into two periods, which we refer to as ‘early’ and ‘late’, that determine the spontaneous and evoked network dynamics (Colonnese and Phillips 2018; Leighton and Lohmann 2016; Luhmann and Khazipov 2018). The division between these epochs occurs before the onset of active sensory experience: birth for auditory, somatosensory and visual systems in humans, eye-opening in the rodent visual system, whisking in rat somatosensory cortex (Colonnese et al. 2010; Fabrizi et al. 2011; Chipaux et al. 2013). These macroscopic changes happen very fast, while many microscopic parameters are observed to vary smoothly during the transition between the early and late periods. In order to show how this can happen while reproducing the experimental data, we start by reviewing some key phenomenological aspects our model aims at reproducing. We then describe an extension of a two-population Wilson-Cowan (WC) model, that allows to parameterize processes related to development (e.g. ratios of the synaptic population onset time delays and of currents received by inhibitory and excitatory cells) which appear to be crucial for the following bifurcation analysis.

### Development of the Thalamocortical loop

This paper will use the rat visual cortex as its reference model as its development has been studied in great detail. Experiments in rat visual cortex show that the switch between early and late periods is extremely rapid, occurring within 12 hours (Colonnese et al. 2010; Colonnese 2014) between P11 and P12. Both visually evoked and spontaneous activity differ qualitatively between early and late period at multiple levels. These include the duration, amplitude and structure of flash evoked activity, the linearity of visual responses, the oscillatory and laminar structure of activity, the continuity of background activity, the stability of membrane potential depolarization, the sparcity of neuronal firing, and the regulation of activity by state (see (Colonnese and Phillips 2018) for review). Some of these differences may be a result of changes in the pattern a activation in the retina. Indeed, retinal activity transitions, at about age 2 postnatal weeks, from intermittent waves of spontaneous activity to persistent, light-regulated activity (Demas et al. 2003), and the processes patterning early retinal activity were recently modeled (Cotterill et al. 2016). Here, we examine the hypothesis that additional changes in the dynamics occurring during the active periods could result from changes in the thalamocortical network, thereby modifying their responses to retinal input.

Most of the changes between early and late periods are attributable to circuit maturation in thalamocortex, however. During the early period, retinal input, which is not oscillatory, drives spindle-burst oscillations in visual thalamus and cortex. This 6-20Hz oscillation, the period of which is age-dependent, synchronize firing within the cortical column (Hanganu et al. 2006; Colonnese and Khazipov 2010; Colonnese et al. 2010; Shen and Colonnese 2016) and between cortex and thalamus (Yang et al. 2016; Murata and Colonnese 2016). These oscillations disappear suddenly at the switch (P12) to be replaced by brief, graded responses to transient retinal activation, and persistent stable depolarization to extended activity (Colonnese et al. 2010; Colonnese 2014). In vivo whole-cell recordings show that during the early stages periods of activity are unstable and the network does not produce an ‘active’ state, defined as a stable depolarized state, continuous during wakefulness, or alternating with down-states during sleep (Fig. 1), until P12. The change to adult-like network dynamics occurs as a rapid switch between P11 and P12 (Colonnese et al. 2010; Colonnese 2014)(Fig. 1). Extracellular recordings no longer reveal a prominent peak in spectral power during wakefulness, also indicating prominent asynchronous state (Harris and Thiele 2011; Shen and Colonnese 2016). Measurements of sensory responsiveness indicate a shift from a non-linear ‘bursting’ regime to a graded, linear regime simultaneous with the switch from unstable/oscillatory to stable membrane currents (Colonnese et al. 2010).

**Figure 1:**
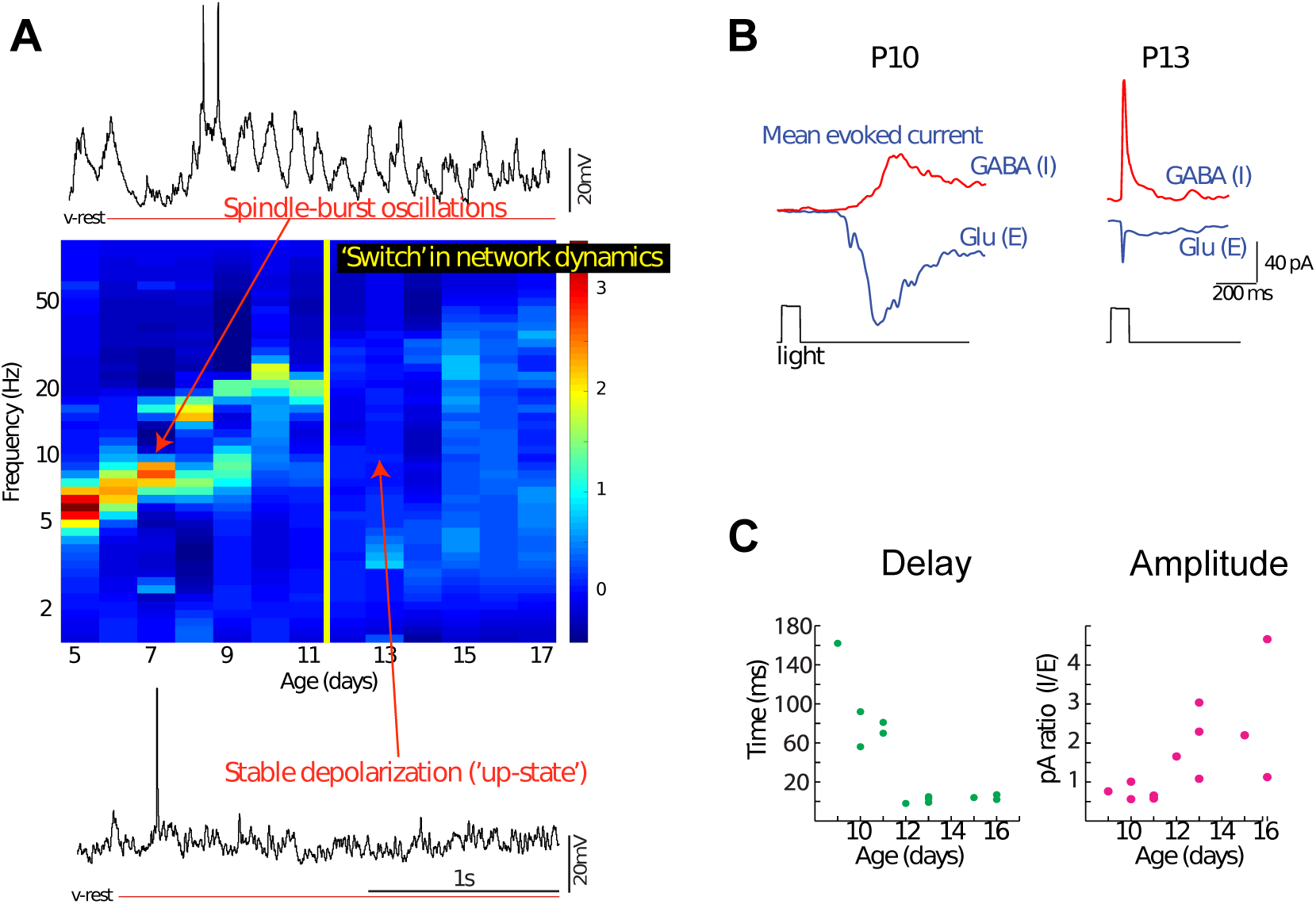
Network dynamics and inhibitory synaptic currents during development in visual thalamocortex (TC). A. Spectral analysis of local field potentials and whole-cell membrane dynamics reveals robust oscillatory behavior in early developing TC that increases in frequency but decreases in amplitude. Transition to an adult-like pattern occurs between P11 and P12. Color graph in the center shows representative spectrograms of layer 4 LFP for each of the indicated ages, with representative whole-cell currents at P7 (top, showing prominent 8Hz oscillations in membrane potential, referred as “spindle-burst” activity) and P13 (bottom, dominated by stable depolarization referred to as an “up” or “active” state). Spectra are normalized to mean power and the 1/f noise is removed by subtraction. Data are novel analysis derived from animals first reported in (Colonnese 2014; Colonnese and Khazipov 2010). B-C. Whole-cell voltage clamp in vivo shows changes in inhibitory currents are strongly correlated with maturation of network dynamics. B. Representative neurons at two ages showing mean light-evoked inhibitory (GABA) and excitatory (Glutamate) currents. Inhibitory currents at young ages are smaller and delayed relative to excitatory currents. C. Development of evoked inhibitory delay (left) and amplitude (right) in visual cortex. Delay is measured as difference between excitatory and inhibitory current onset, and amplitude as the ratio between the peaks of each current. Data are reproduced from (Colonnese 2014).

Together these observations show that the first two weeks of the rat’s brain development are characterized by a thalamocortical network that maintains an oscillatory activity. The maturation of these early dynamics to adult-like networks occurs as a sudden transition in the network dynamics leading to an adult-like linear processing regime, asynchronous network activity and stable depolarization during activation. As measured in somatosensory and visual cortex, the synaptic changes most closely associated with this developmental switch are an increase in the amplitude of synaptic inhibition and a decrease in its delay (Minlebaev et al. 2011; Colonnese 2014). It appears that through this process inhibition and excitation drive towards a general balance. Such measurements of feed-forward inhibition in cortex suggest that changes in the timing and strength of inhibition are important, but they reflect only a limited population of total inhibition that likely evolves more gradually. We therefore take these limited measurements of inhibitory and excitatory synaptic currents as starting point to examine timing and power of inhibition during thalamocortical network development.

### The mathematical model

To clarify the potential key role of inhibitory maturation in the developmental switch, we developed a firing rate model explicitly accounting for the synaptic timing effects To this end, we extended the WC model in order to allow us to study independently the effects of excitatory and inhibitory synaptic delays, as well as their rise and decay times (Fig. 2A). Evoked inhibitory and excitatory synaptic population currents seen experimentally (Fig. 1B) display double exponential profiles not accounted for by the instantaneous exponentially decaying currents of the classical WC model. These are well parameterized by second-order dynamical systems with double-exponential impulse responses, whose amplitude, rise and decay times can be prescribed (see Fig. 2).

**Figure 2:**
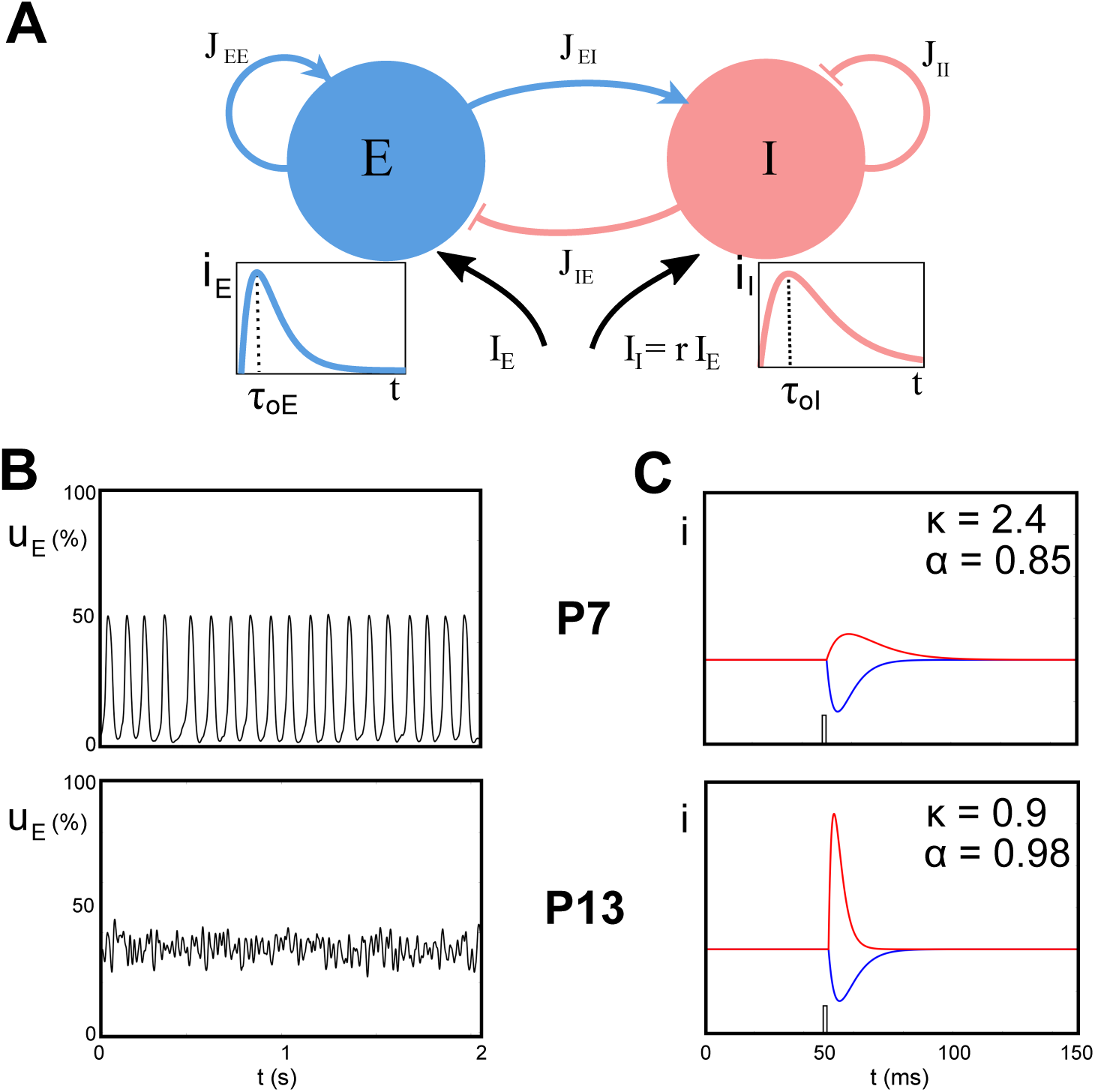
Two-populations model with double exponential synapses. (A) Two populations of neurons, one excitatory (blue) and one inhibitory (red) in interaction (coupling coefficients *J*_*EE*_, *J*_*EI*_, *J*_*IE*_, *J*_*II*_) and receiving external inputs, respectively *I*_*E*_ and *I*_*I*_. Insets display the typical double exponential responses to impulses for each population (arbitrary units), with identical peak amplitudes and respective onset delays *τ*_*oE*_ and *τ*_*oI*_. (B) Numerical simulations of the model, with Gaussian noise stimulus, at P7 (top) and P13 (bottom). In these graphs is represented the proportion of active excitatory neurons as function of time: at P7, the network displays an oscillatory behavior at around 10Hz; while at P13, the activity shows a noisy stationary behavior, both being comparable to the *in vivo* recordings (Fig. 1). (C) Excitatory (blue) and inhibitory (red) synaptic currents in response to a brief impulse, in arbitrary units, with the convention that the total excitatory transmitted current (area under the curve) is normalized to 1.

In detail, synaptic responses are defined as the solutions of:

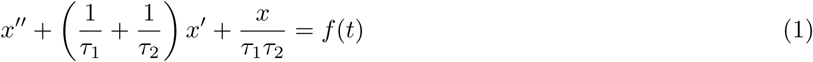

where *x*′ (resp. *x*′′) denotes the first (resp. second) derivative in time of *x* and *f* (*t*) the input to the system. We study in more detail these equations in the SI text and recall that the solution is given by a difference of two exponentials with rates having a ratio *λ* = *τ*_2_*/τ*_1_, thus *τ*_1_ can be interpreted as a rise time and *τ*_2_ as a decay time assuming *τ*_1_ > *τ*_2_. Following the approach introduced by Wilson and Cowan (Wilson and Cowan 1972), we model (*u*_*E*_, *u*_*I*_) as the fractions of cells in the excitatory (E) and inhibitory (I) populations firing at time *t*. Each population is assumed to satisfy an equation of type (1), with respective rise and decay times 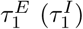 and 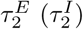, yielding the system of differential equations:

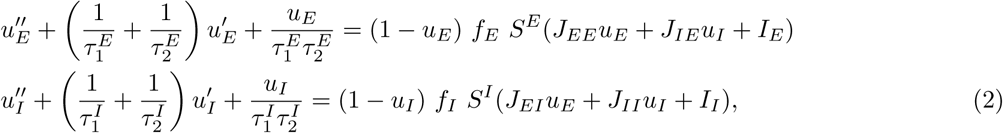

where the input to population E/I possibly induces an activation to the fraction of quiescent cells (1 − *u*_*E/I*_) with a maximal rate *f*_*E/I*_, and according to a sigmoidal transform *S*^*E/I*^ of the total current received. The sigmoids associated to each population may have distinct thresholds *θ*_*E/I*_ and distinct slopes *a*_*E/I*_, but are assumed to have the same functional form: *S*^*E*^(*·*) = *S*(*a*_*E*_, *θ*_*E*_, *·*) and *S*^*I*^ (*·*) = *S*(*a*_*I*_, *θ*_*I*_, *·*) with

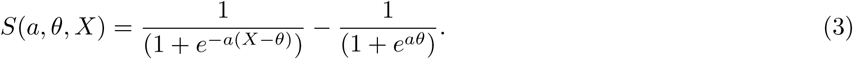

The total input received by a population *E/I* is given by the sum of the external current *I*_*E/I*_ and the input of other cells, proportional to the fraction of firing cells in both populations, and weighted by synaptic coefficients (*J*_*ij*_)_(*i,j*)∈*{E,I}*_, that depend both on the level of connectivity between populations (*i, j*) and on the average amplitude of the post-synaptic currents^1^.

To reduce dimensionality we now express time in units of 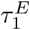 (dimensionless time 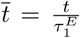), yielding a system depending on three dimensionless ratios:

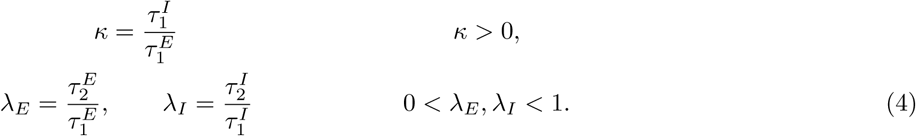

The coefficients *λ*_*E*_ and *λ*_*I*_ determine the slope of the decay of the double-exponential synaptic responses, while *κ* provides a proxy for the ratio between inhibitory/excitatory currents onset delay ^2^.

These two parameters are key for the problem at hand. Indeed, experimental data (Fig. 1) show a decrease of the inhibition onset delay relative to the excitatory one, hence we expect *κ* to decrease during development.

Moreover, the system depends upon two amplitude parameters *f*_*E*_ and *f*_*I*_. Timescales of currents not only impact the timing of synaptic rise and decay; they also affect the total current transmitted, that we can quantify as the integral of the impulse synaptic response (response to a Dirac input), denoted *A*_*E*_ and *A*_*I*_. We fixed *f*_*E*_*λ*_*E*_ = 1 to satisfy the normalization *A*_*E*_ = 1, and denoted by *α* the ratio *A*_*I*_ */A*_*E*_ (see SI Text):

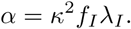

Heuristically, when *α* < 1, the total current that would be received by the inhibitory population in response to a pulse of current is smaller than the total current received by the excitatory population in the same situation. In terms of this interpretable parameter, the equations (2) reduce to:

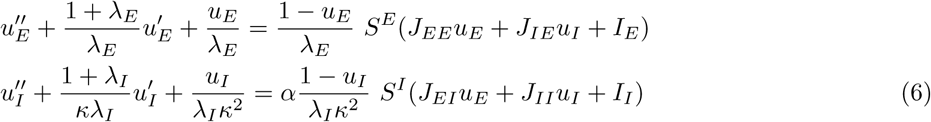

where the derivatives are now taken with respect to the dimensionless time variable 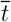.

To recapitulate, in our reduced model, there are two key quantities that we will work with: *κ* that gives the ratio of the relative onset-delay of inhibition and excitation and *α* giving the ratio of the total excitatory and inhibitory transmitted currents. In other words, heuristically, *κ* indicates the temporal scale of inhibition relative to excitation and *α* is the relative total impact strength of inhibition vs. the excitation. These two control the balance between excitation and inhibition in the model.

### Mathematical and Computational methods

The full mathematical framework is provided in the reference (Wilson and Cowan 1972) and in the main text, while more details can be found in SI text. The bifurcation analysis have been performed using XPPAUT (Ermentrout 2002) and Matcont (in Matlab environment) (Dhooge et al. 2003). The results shown in Fig. 2B, Fig. 5, Fig. 6 and Fig. 7, are based on numerical simulation where the Eqs. 6 have been implemented in Matlab and Python codes with the Euler method. In particular, for each set of parameters, we simulated 1 second of dynamics, while frequencies (by FFT analysis) and amplitudes of the oscillations have been calculated only on the last 500 ms, to avoid the initial transient. Moreover, for the cases with noise, for each set of parameters we run 10 different simulations, and shown the mean for the amplitudes of the oscillations in Fig. 7.

**Figure 3:**
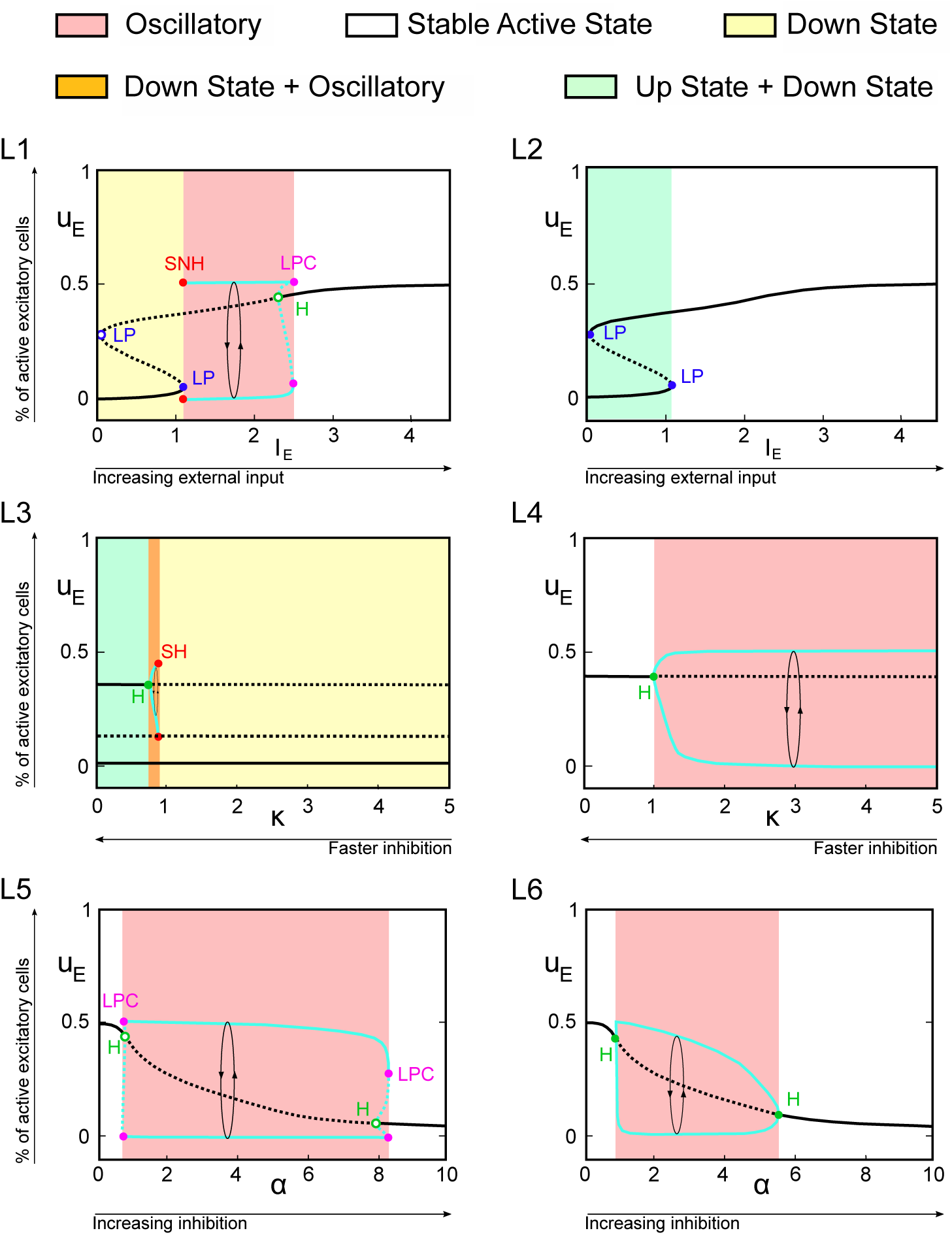
Bifurcation diagrams in 1-dimensional parameter spaces: In all panels we show the value of the excitatory activity *u*_*E*_ associated with equilibria and periodic orbits (in that case, maximal and minimal values are plotted) as a function of one given parameter. L1-L2 show the influence of *I*_*E*_ for fixed *α* = 1 and *κ* = 3 (L1) or *κ* = 0.5 (L2). L3-L4 elucidate the influence of *κ* with *α* = 1 and *I*_*E*_ = 0.75 (L3) or *I*_*E*_ = 1.5 (L4). L5-L6 characterize the influence of *α*, with *I*_*E*_ = 1.5 and *κ* = 4 (L5) or *κ* = 1.5 (L6). Regions are colored according to the type of stable solutions. We distinguish between up and down states when more than one solution is present for a given choice of the parameters, while we call it generically stable active state, when only one (stable) constant solution is present. Bifurcation labels: **LP**: Limit point, **H**: Hopf, **SH**: Saddle homoclinic, **SNH**: Saddle node homoclinic, **LPC**: Limit point of cycles. Solid (dashed) black lines represent stable (unstable) solutions for *u*_*E*_. Maximal and minimal *u*_*E*_ along cycles are depicted in cyan (solid: stable, dashed: unstable limit cycles).

**Figure 4:**
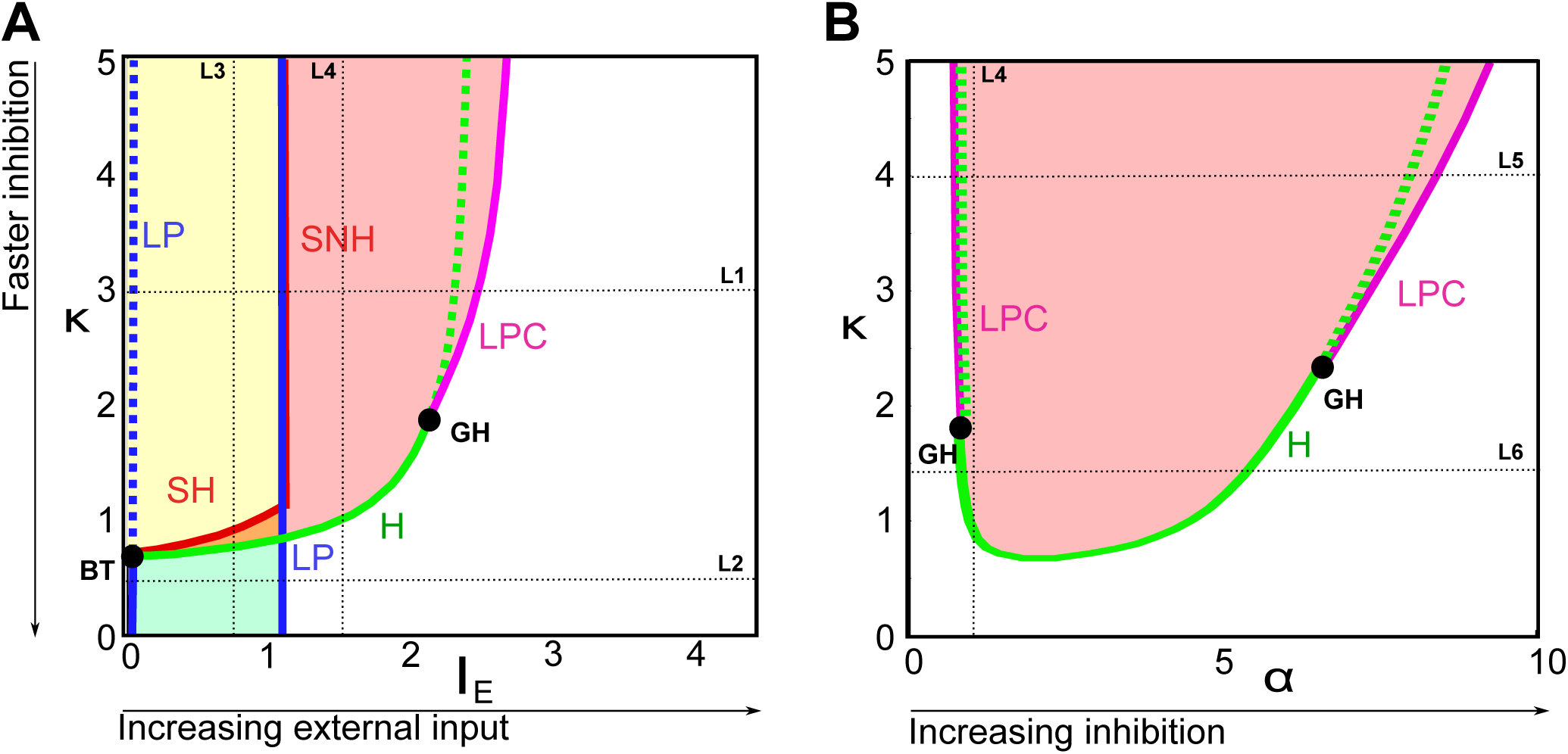
Codimension-two bifurcation diagrams: (A) elucidates the transitions in the network behavior as a function of input and inhibitory timescales (*I*_*E*_, *κ*) for fixed *α* = 1; (B) characterizes the network behavior as a function of amplitude and timescale of inhibition (*α, κ*), for fixed *I*_*E*_ = 1.5. Color-code and notations as in Fig. 3. Codimension-two bifurcations: **BT**: Bogdanov-Takens, **GH**: Generalized Hopf (Bautin). Solid and dotted blue (resp. green) lines represents respectively stable or unstable limit points (resp. Hopf bifurcations). Sections corresponding to the different panels of Fig. 3 are depicted with dotted black lines.

**Figure 5:**
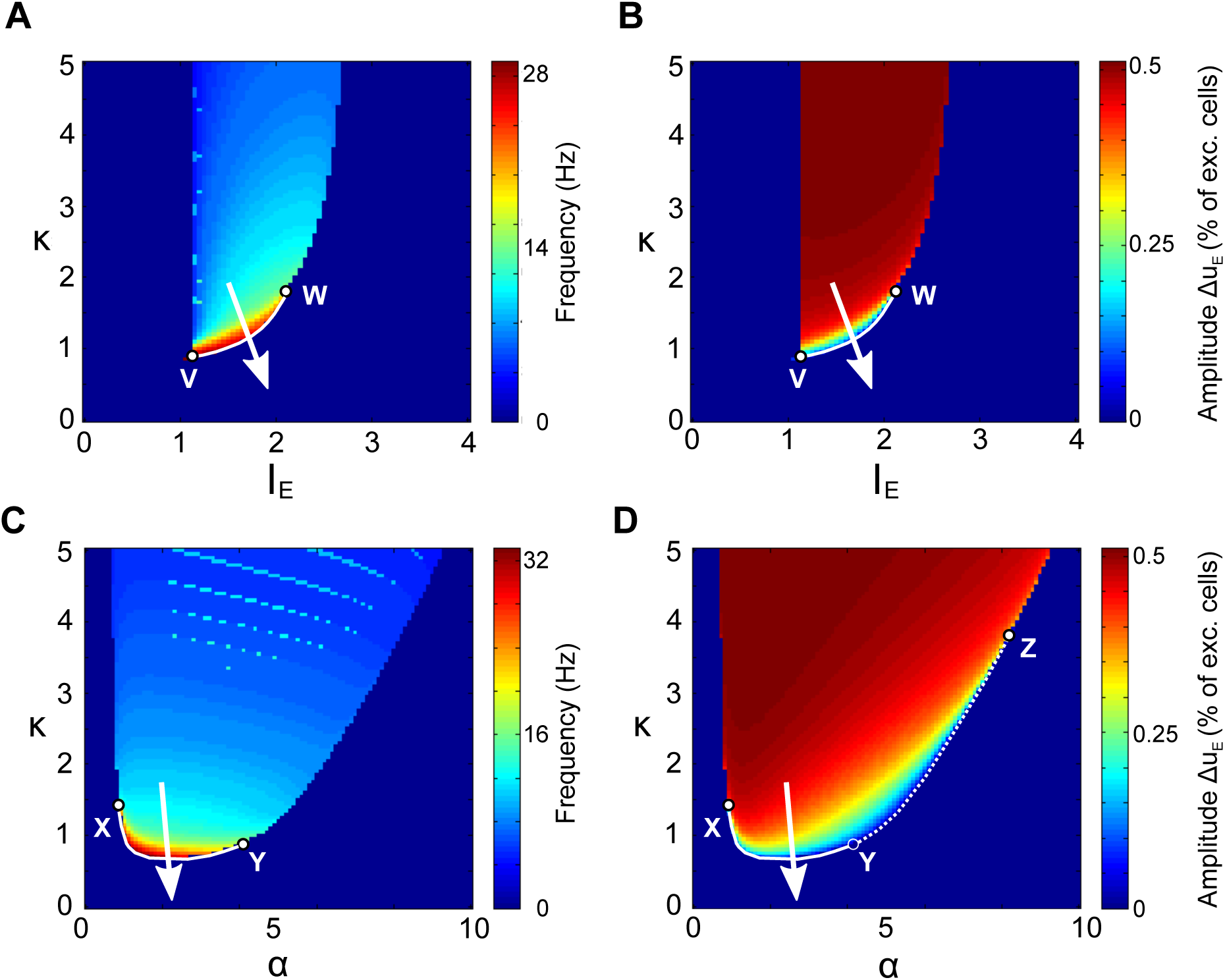
Frequency and amplitude of the oscillatory solutions. Panels A and B refer to (*I*_*E*_, *κ*) phase space with *α* = 1, while panels C and D to (*α, κ*) phase space with *I*_*E*_ = 1.5. In panels A and C the frequency is expressed in Hz, after fixing 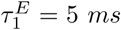 (thus, the excitatory onset delay has the reasonable value of *τ*_*oE*_ ∼ 4.5 ms). Suitable trajectories during development should cross the white line between the points V and W at the moment of the switch. In panels B and D the amplitude values correspond to the difference between the maximum and the minimum of the oscillation in the excitatory population activity. Suitable trajectories during development should cross the white line between the points X and Y at the moment of the switch, while the dotted white line between Y and Z would only allow for a decrease in the oscillations amplitude.

**Figure 6:**
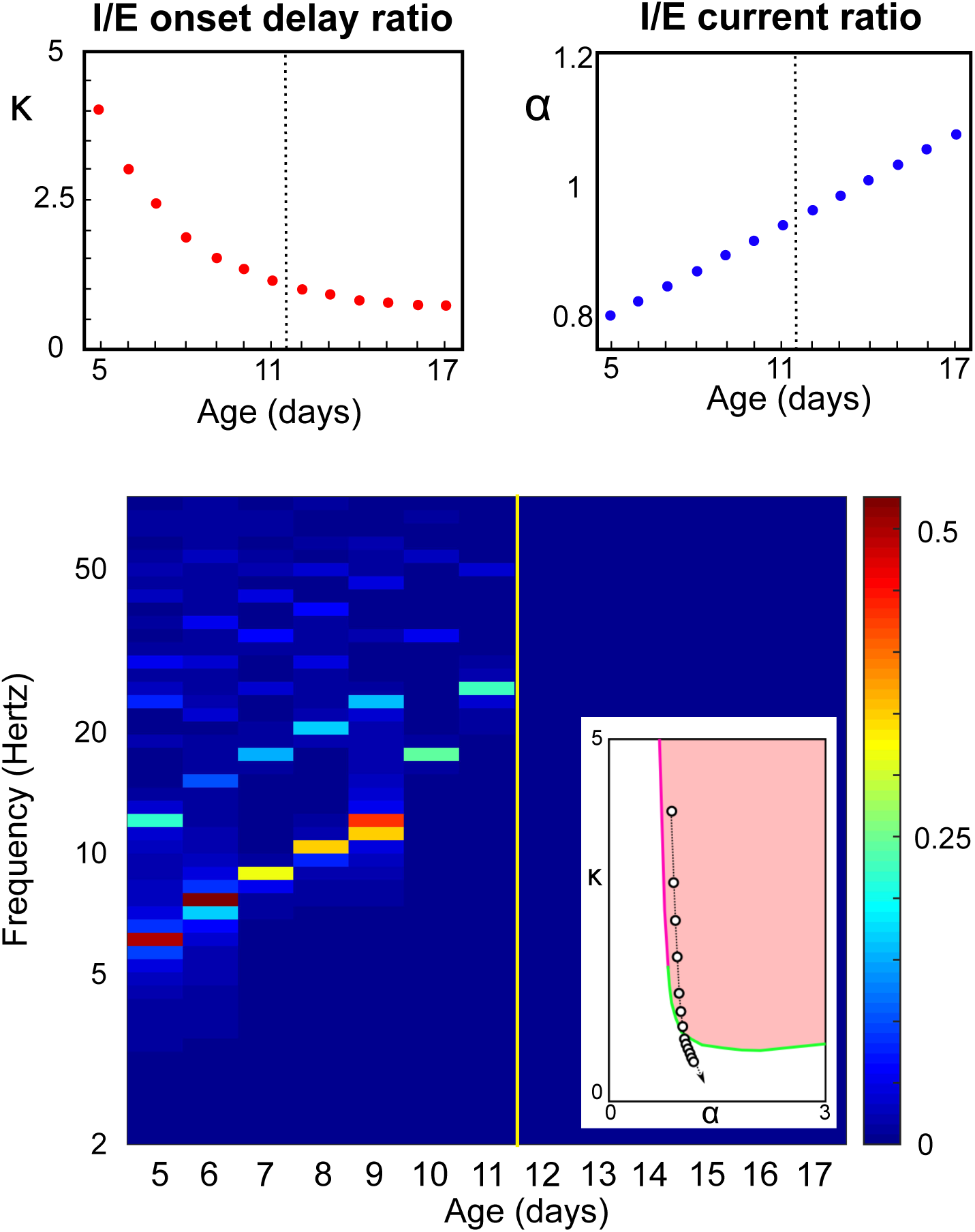
Model of the developmental switch. Top: Proposed developmental trajectories for relative inhibitory onset delay *κ* and amplitude *α*. The black dashed vertical lines indicates the moment of the development switch, happening between P11 and P12. Bottom: Spectrogram of the solutions of Eq. 6 with *I*_*E*_ = 1.5 and 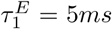, computed as the fast-Fourier transform of the excitatory activity *u*_*E*_; same calculation and representation as used in the experimental data of Fig. 1(A). In the inset, the chosen trajectory in the bifurcation diagram in the 2-dimensional parameter space (*α, κ*). Color codes as in Fig. 3 and Fig. 4.

**Figure 7:**
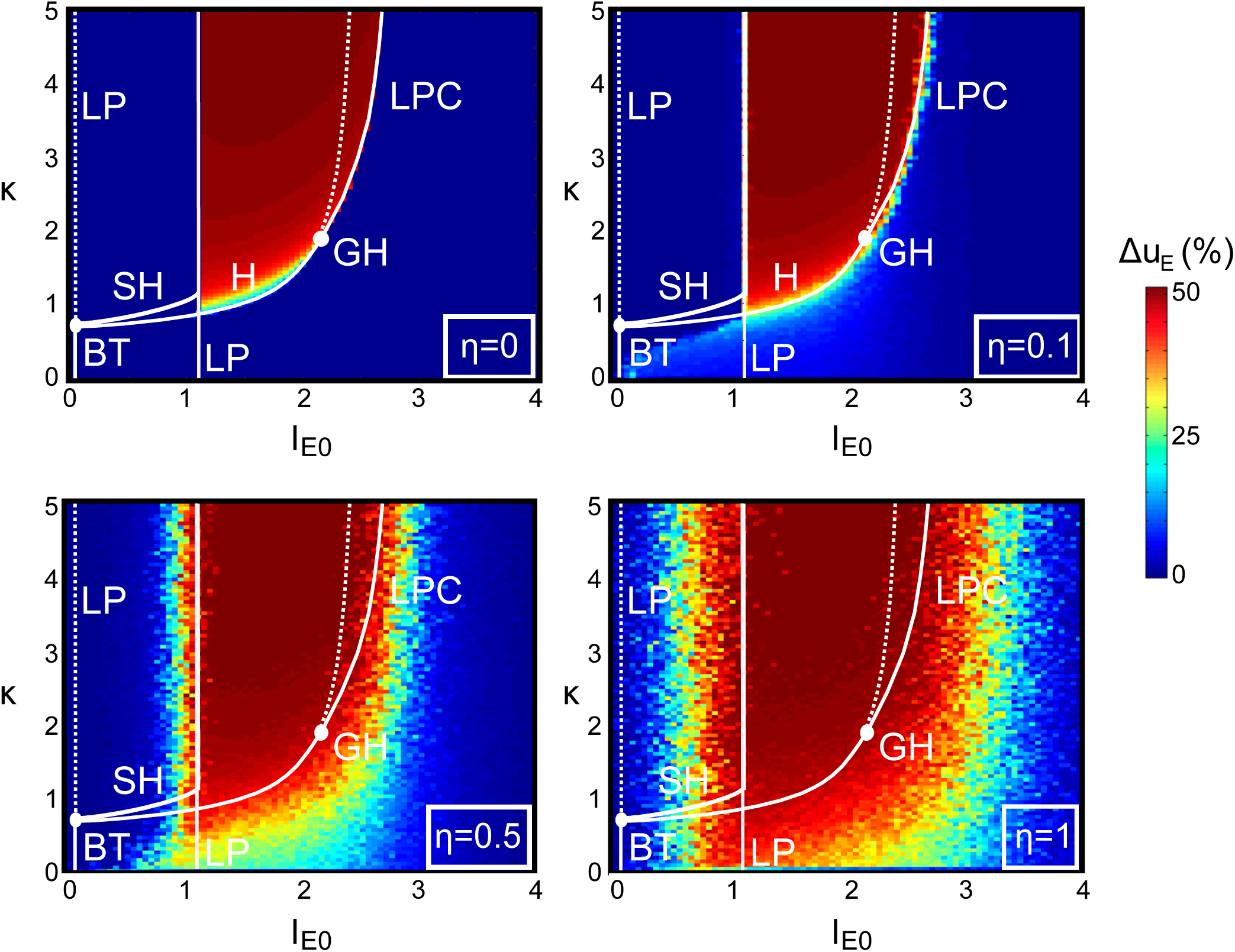
Amplitude of the oscillatory solutions in the presence of noise. We consider the effects of Gaussian noise on the statistics of Fig. 5, namely the difference between the maximum and the minimum values of the excitatory activity *u*_*E*_, in the (*I*_*E*0_, *κ*) phase space. The codimension 1 and 2 bifurcations of the noiseless case are indicated, following the same notation as in the Fig. 4.

## Results

The patterns of activity recorded experimentally through the switch reveal multiple aspects that a model should reproduce in their spontaneous activity:

(c1) a smooth increase in spontaneous oscillation frequency;

(c2) a relatively narrow oscillatory band from theta to low gamma;

(c3) a gradual decrease in oscillation amplitude, and

(c4) a sudden transition to broad-band high gamma oscillations (asynchronous Up-state).

Moreover, responses to brief stimuli also significantly change during development, a feature that a model accurately accounting for in vivo development should also reproduce:

(c5) a transition from a highly non-linear all-or-none response to transient inputs to a linear graded response reflecting the stimulus amplitude and duration,

(c6) this transition occurs at the same time as the phase transition observed for spontaneous activity.

We thus investigate the spontaneous and driven properties of this system as a function of the parameters altered during development, in order to assess whether – and under which conditions– the model can reproduce experimental dynamics, and what these conditions tell us about the underlying biological system.

### Spontaneous dynamics

We thus characterized the attractors of system 6 and their stability (steady-states or periodic orbits) as a function of the parameters altered during development. Our assumption is that stable attractors determine the observable dynamics in our network and these should correspond to developmental epochs defined by consistent activity patterns (e.g, the early period of oscillations vs later period of bistability), and change qualitatively at bifurcation points that should correspond to the rapid switches observed in cortical activity. Three key parameters are significantly modified during development: *κ* (onset delay of inhibition relative to excitation), *α* (ratio of inhibitory vs excitatory impulse current) and *I*_*E*_ (level of external input to both populations).

Other parameters were set as in standard references (Wilson and Cowan 1972). In particular, we fixed *a*_*E*_ = 1.3, *θ*_*E*_ = 4, *a*_*I*_ = 2 and *θ*_*I*_ = 3.7. The connectivities have been fixed to *J*_*EE*_ = 16, *J*_*II*_ = −3, *J*_*EI*_ = −*J*_*IE*_ = 10. For simplicity, we consider *I*_*I*_ = *rI*_*E*_ with *r* = 0.5 fixed and *λ*_*E*_ = *λ*_*I*_ = 0.8. In the SI text we show that other choices of *r, λ*_*E*_, *λ*_*I*_ and *J* do not qualitatively affect the dynamics of the system in the (*α, κ, I*_*E*_) space.

#### Strong influence of synaptic transmission on network dynamics

The relative timescales and amplitudes of currents transmitted control tightly the dynamics and modify substantially the response of the system to an input. Indeed, Fig. 3 shows the bifurcation diagram of the system for various choices of synaptic parameters. We find that, consistent with the experimental observation, our system features:

1. Stationary states with possible multi-stability between: an *up state* of high E and I activity evocative of the awake adult activity, and a *down state*, associated with low activity akin to the inactive periods during early development, green regions in Fig. 3. or single stable equilibria (referred to as *stable active state* in the sequel irrespective of their activity levels, white parameter space in Fig. 3).
2. Oscillatory states where the E/I populations are activated rhythmically (pink region in Fig 3), reproducing to the periods of activation during early development, and observed for intermediate values of input or inhibition amplitude (panels L1, L5 and L6) or slow inhibition (panel L4), possibly co-existing with the presence of a stable down-state (orange region in Fig. 3).

The boundaries between regions with prescribed numbers of attractors of a given stability are given by the bifurcations of the systems (circles in Fig. 3). We found the following bifurcations:

- Appearance or disappearance of stationary states are associated with *saddle-node bifurcations* (or Limit Points, LP, blue circles Fig. 3). We distinguish two types of saddle-node bifurcations: those yielding one stable and one saddle equilibria (circles filled in blue), and those yielding two unstable equilibria (circles filled in white, associated with the emergence of one repelling and one saddle equilibria)
- Emergence of oscillatory behaviors is related to the presence of *Hopf bifurcations* (green circles labeled H in Fig. 3). Locally, Hopf bifurcations give rise to periodic orbits that are either stable or not and we have identified each case (super- or sub-critical Hopf bifurcations are depicted with green- or white-filled circles respectively).
- The periodic orbits also undergo bifurcations that affect network behavior: they may disappear through homoclinic bifurcations (red points, labeled SNH when corresponding to Saddle-Node Homoclinic, of SH when corresponding to Saddle-Homoclinic) at which point of branch of periodic orbit collides with a branch of fixed points (either a saddle-node point or a saddle equilibrium), and subsequently disappears. We also observed the presence of folds of limit cycles (or limit points of cycles, LPC, in magenta), particularly important in the presence of sub-critical Hopf bifurcations since they result in the presence of stable, large amplitude cycle.

In addition to these observations, Fig. 3 highlights the fact that the location and the mere presence of the above-described bifurcations may change as a function of the other parameters. We elucidate these dependences in Fig. 4 by computing the codimension-two bifurcation diagrams of the system as inhibitory timescale and external current (*κ* and *I*_*E*_, panel A) or inhibitory timescale and inhibitory currents amplitude (*κ* and *α*, panel B) are varied together. In these codimension-two diagrams, because of the structural stability of saddle-node and Hopf bifurcations, reveal curves of saddle-node (blue) and Hopf (green) bifurcations, with singularities located at isolated *codimension-two* bifurcation points. Codimension-one diagrams depicted in Fig. 3 correspond to sections of codimension-two diagrams, and we indicated the location of these sections in Fig. 4. In both panels, we observe the presence of two codimension-two bifurcation points:

- A Bogdanov-Takens bifurcation (BT Fig. 4(A)). At this point, the Hopf bifurcation meets tangentially the saddle-node bifurcation curve and disappears, explaining in particular the transition between panels L1 (no Hopf bifurcation, on one side of the BT bifurcation) and L2 (on the other side of the BT bifurcation). The universal unfolding of the Bogdanov-Takens bifurcation is associated with the presence of a saddle-homoclinic bifurcation, SH curve, along which the periodic orbit disappears colliding with a saddle fixed point. We have computed numerically this curve (depicted in red in Fig. 4), and found that this curve collides with the saddle-node bifurcation associated with the emergence of down-states, at which point it turns into a saddle-node homoclinic bifurcation (namely, instead of colliding with a saddle fixed point, the family of periodic orbits now collides with a saddle-node bifurcation point). From the point of view of the observed behaviors, this transition has no visible impact. This curve accounts for the presence of SNH and SH points in Fig. 3, panels L1 and L3.
- Bautin bifurcation points (or Generalized Hopf, labeled GH), where the Hopf bifurcation switches from super- to sub-critical, underpinning the variety of types of Hopf bifurcations observed in Fig. 3, particularly between panels L5 and L6. The universal unfolding of the Bautin bifurcation is associated with the presence of a fold of limit cycles (or Limit Point of Cycles, LPC, depicted in magenta), that was already identified in panels L1 and L5 and which appear clearly, in Fig. 4, to be those sections intersecting the LPC lines.

### Model validation and interpretation

#### Phase transition

The bifurcation diagrams point towards parameter regimes exhibiting dynamics compatible with the spontaneous activity observed experimentally in vivo at different stages of development. Furthermore, this analysis suggests how parameters may vary simultaneously for recovering the developmental switch.

In detail, the oscillatory activity characteristic of immature animals (P5-10) can be found in both orange and pink regions of Fig. 3 and Fig. 4, and asynchronous activity characteristic of late development (P≥12) corresponds to up-states of bistable (green) parameter regions or to parameters characterized by a unique active state (white regions). Any trajectory in the parameter space connecting these two different scenarios (orange or pink towards green or white) would therefore allow a phase transition satisfying the condition (c4) listed above.

In terms of biophysical parameters, oscillations are particularly prominent for *κ* sufficiently large, i.e. when inhibitory currents activate after a delay that is larger than the excitatory currents (excitation leads inhibition), and only for intermediate values of input *I*_*E*_ (too low or too high input leading to down- or the up-states, L1 of Fig. 3). Low values of *κ*, associated with faster inhibitory currents, yield dynamics generally consistent to later developmental stages devoid of intrinsic oscillations.

When *κ* is progressively varied for a fixed value of the input below the saddle-node bifurcation (Fig. 3-L3), the dynamics consistently display bistability between up-states and down states for low *κ*, with an up-state losing stability through a supercritical Hopf bifurcation, yielding a narrow oscillatory regime as *κ* is further increased, disappearing through saddle-homoclinic bifurcation, and leaving the system in down-state. For larger *I*_*E*_ (Fig. 3-L3), small *κ* regimes are associated with up-states losing stability through a super-critical Hopf bifurcation as *κ* is increased (for *I*_*E*_ smaller than the value associated to the codimension-two Bautin bifurcation observed in Fig. 4(A), sub-critical otherwise), and leading to periodic dynamics.

In total our analysis indicates that as the inhibition onset speeds-up, the network dynamics pass through bifurcations that bring it from an oscillatory regime with relatively low oscillation frequencies and large amplitudes to a regime with a stable active constant activity state, that we can interpret as an up-state. Indeed, as we can see in the codimension-two bifurcation diagrams Fig 4(A), for any given value of the external input *I*_*E*_, there exists a critical value *κ*_*c*_(*I*_*E*_) at which the system undergoes a bifurcation towards either a unique up-state regime (white region) or a bistable regime (green region). A similar transition is observed experimentally around P11-P12, characterized by a switch from an oscillatory to a prominent up-state of activity. For *κ* < *κ*_*c*_(*I*_*E*_), the bifurcation diagrams Fig. 4(A) and Fig. 3(L2) show indeed that the system tends to settle into the active stationary state regime (pink towards white regions), except for small values of input *I*_*E*_ for which this up-state may co-exist with a down-state (orange towards green regions). Interestingly, we notice that the value of the input *I*_*E*_ associated with the saddle-node bifurcation (blue line in Fig.4) is independent of *κ*, indicating that the emergence of the down-state arises at levels of thalamic or cortical input that are independent of the inhibitory/excitatory current onset delay ratio. In other words, the down-state may be observed at any stage of inhibitory maturation if the feedforward input is sufficiently low.

In vivo (Fig. 1)the amplitude of inhibitory synaptic currents increase relative to the excitatory currents (parameterized by *α* in the model) during development. In the model, we observed (Fig. 4(B)) that, for input levels associated with oscillations at *α* = 1, the curve of Hopf bifurcations in the (*α, κ*) plane is convex, so that increasing or decreasing the relative amplitude of inhibitory currents *α*, or simply decreasing the relative inhibitory onset delay *κ* beyond a critical value, yields a switch to non-oscillatory regimes similar to the adult network responses. We note that, although the interval of values of *α* associated with oscillatory behaviors shrinks as *κ* decreases (as clearly visible in Fig. 3 (L5, L6)), the choice of *α* = 1 to compute Fig. 4(A) is generic, as any value between about 1 and 7 would have given qualitatively identical results.

We further note that for *κ* large, the Hopf bifurcations are sub-critical. While this does not affect the presence of periodic orbits (because of the limit points of cycles evidenced), it will affect the finer structure of how the network transitions to oscillations. In that regime, contrasting with the progressive disappearance of oscillations typical of super-critical Hopf bifurcations, we would observe that large-amplitude oscillations disappear suddenly as they reach the limit point of cycles. Moreover, near this region, we may have a coexistence between the stable oscillations and stable (up- or down-) constant states, which, in the presence of noise, may generate inverse stochastic resonance.

#### Co-variation of relative synaptic delays and amplitudes and developmental switch

The physiological situation with gradual smooth variations in synaptic parameters leads to a sudden and dramatic switch in the network dynamics when these parameters cross one of the bifurcations observed. In particular, three possible scenarios can lead the system to transition from rhythmic to stationary activity consistent with the experimental evidence: increase of the input, decrease of the inhibitory onset delay relative to the excitatory one (*κ*), increase of inhibitory current amplitude relative to the excitatory one (*α*). To determine which transition is more consistent with the physiological measurements, we now consider whether these scenarios are consistent with the characteristic increase in oscillation frequency observed experimentally (criterion c1, see Fig. 1(A)).

A robust measurement of this acceleration was obtained by deriving the power-spectrum of the recorded activity. To directly compare model dynamics to the experimental measurements, we compute the fast Fourier transform of the solutions to Eq. 6 within the range of parameters covered by the above bifurcation analysis, and report in Fig. 5 the two relevant quantities associated: peak frequency and amplitude. In these diagrams, the region of oscillations, delineated in Fig. 4 by the Hopf bifurcation and limit points of cycles, clearly stands out. Rhythmic immature activity suggests that associated parameters lie within the oscillatory region (pink in Fig. 4), and leave this region as development progresses. Fourier analysis of the solutions reveals both an increase in oscillation frequency – condition (c1) above – and a smooth decrease in oscillation amplitude – condition (c3) – in the vicinity of the supercritical Hopf bifurcation, particularly along the white curves depicted in in Fig. 5A (linking points V and W) and Fig. 5C (between the points X and Y). Both curves show a relatively small variation in *κ*, suggesting that the onset delay of inhibition plays a central role during the developmental switch. While changes in *I*_*E*_ or *α* theoretically could underpin a sudden switch between an oscillatory state and an constant activity state, the telltale frequency acceleration near the transition specifically points towards a decrease of the relative inhibitory current onset delay *κ*. In detail, a constant or increasing *κ* but with increasing *α* would imply a smooth decrease in oscillations amplitude, when crossing the white line of Fig. 5D between the points Y and Z. However, this trajectory would not produce a simultaneous acceleration in the oscillation frequency.

Notice that since in the phase space, the oscillation frequency increase happens only near the Hopf bifurcation line, our model predicts that the corresponding oscillatory amplitude should progressively decrease, and not suddenly drop as would be the case if the switch occurred by crossing the Limit Point of Cycles (LPC) or the homoclinic bifurcation. In other words, condition (c3) arises naturally in the model when conditions (c1) and (c4) are satisfied.

Following our analysis we can now propose a specific developmental variation of the parameters, or a set of parametric paths, that reproduces accurately changes in both the macroscopic dynamics and the spectrogram seen in the experiments (see Fig.1C). In order to follow the observed data, we constrained qualitatively the inhibitory onset timescale to decrease and the synaptic amplitude to increase with the developmental stage as seen in data. We fixed the excitatory time constant to a realistic value of 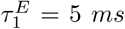. A specific set of age-dependent parametric trajectories depicted in Fig. 6 (top), resulted in an accurate match with the experimental data: a sudden switch, at age P12, between an oscillatory state and a quasi-constant activity, together with a clear increase in oscillations frequency approaching the switch.

#### Robustness to noise

The above results were obtained in a deterministic system, which allowed us to use bifurcation theory to show sudden transitions in the network activity as a function of the excitation/inhibition time-scale ratio and input level. However, in physiological situations, neural populations show highly fluctuating activity due to multiple sources of noise (e.g. channel noise or synaptic noise resulting from the intense bombardment of neurons from other brain areas) (Faisal et al. 2008). The role of noise in dynamical systems has been the topic of intense study (Lindner et al. 2004; Berglund and Gentz 2006). In general, large amounts of noise generally overwhelm the dynamics, while small noise produces weak perturbations of the deterministic trajectories away from instabilities. In the vicinity of bifurcations, even small amounts of noise may significantly modify the deterministic dynamics (Lindner et al. 2004; Tuckwell et al. 2009; Buchin et al. 2016). Because of the importance of bifurcations in the switch, we investigate in this section the role of noise on the dynamics of Eq. 6. We assume that the various sources of noise result in random fluctuations of the current received by each population around their average value *I*_*E*0_ and 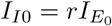, and model the current received by each population by the Ornstein Uhlenbeck processes:

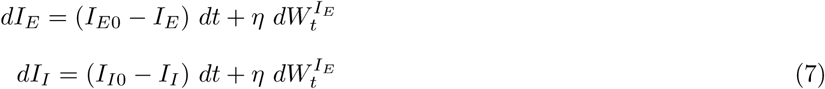

where 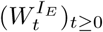 and 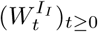 are standard Wiener processes and *η* is a non-negative dimensionless parameter quantifying the level of noise. We computed the solutions of the system and derived the statistics of the solutions for various level of noise. The average amplitude of the variations in time of the solutions as a function of the excitatory input *I*_*E*0_ and excitation/inhibition timescale ratio *κ* is depicted in Figure 7 for various values of *η* and fixed *r* = 0.5. Away from the bifurcation lines, the impact of moderate noise is reasonably negligible, and the noisy system behaves as its deterministic counterpart, as we can see in Fig. 2B, where at P7 (before the switch) the oscillating regime displays minor perturbations from the deterministic (*η* = 0.3, *κ* = 2.4, *α* = 0.85, *I*_*E*_ = 1.5), and similarly the stationary up-state at P13 (after the switch) is only perturbed (*η* = 0.3 *κ* = 0.9, *α* = 0.98, *I*_*E*_ = 1.5). Near transitions, we observe two effects that have a moderate, yet visible effect, on the precise parameters associated to a switch. In particular, we observe that all transition lines tend to be shifted by the presence of noise, enlarging the parameter region associated with oscillations. This effect is likely due to stochastic resonance effects in the vicinity of heteroclinic cycles or folds of limit cycles, while in the vicinity of the supercritical Hopf bifurcation, coherence resonance effects due to the interaction of noise with the complex eigenvalues of that equilibrium (see (Lindner et al. 2004)).

We thus conclude that despite a slight quantitative shift of the parameter values associated with the switch, the qualitative behavior of the noisy system remains fully consistent with the experimental observations. Indeed, a decrease in the timescale ratio *κ*, possibly associated with an increase in the input, yields a transition from oscillating solutions to a stationary activity level associated with linear responses to input, which ensures that the proposed mechanism underlying the switch is robust in the presence of noise.

#### Response to transient inputs

In the previous section we have shown under which constraints all conditions (c1)-(c4) are satisfied for the model to match the experimental data on the spontaneous activity during the development. However, spontaneous activity is not the only dramatic change reported during the developmental switch: the responses to transient input, like light pulses, are also markedly modified through the switch. These phenomena, labeled conditions (c5) and (c6) above, can be seen as a consequence of the same bifurcations arising in the system. Having developed and identified the parametric paths in the model solely for the spontaneous activity, we wanted to see if the model and our chosen trajectory through *κ, α* and *I*_*E*_ also generate realistic responses to stimuli. We thus considered the dynamics of the system at P7 (*κ* = 2.4, *α* = 0.85, *I*_*E*_ = 1.5) and P13 (*κ* = 0.9, *α* = 0.98, *I*_*E*_ = 1.5) in response a brief square pulses (duration *δt* = 10ms) of varying amplitude Δ*I*_*E*_ emulating light pulses:

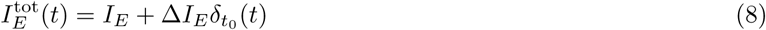

where 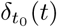 is equal to 1 for *t* ∈ [*t*_0_, *t*_0_ + *δt*] and 0 otherwise.

At P7, the pulse causes both a transient increase in the amplitude of the excitatory activity, and, more markedly, a phase-shift of the spontaneous oscillations, that depends on the phase at which the stimulus is applied (Fig. 8A). The amplitude of the transient response does not grow linearly with the amplitude Δ*I*_*E*_ of the external transient. In sharp contrast, after the switch, the response to the pulse is characterized by small-amplitude damped oscillations whose maximal amplitude shows a much more gradual dependence in the pulse amplitude, and no specific dependence on the timing of the impulse presentation (Fig. 8A, P13). To assess, before the switch, the dependence of the response to the stimulus, we considered the phase shift Φ_*t*_, implicitly defined asymptotically by

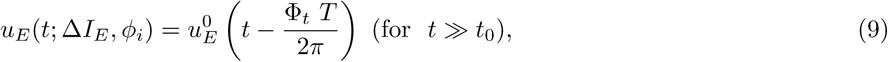

where 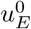 is the excitatory activity for Δ*I*_*E*_ = 0, *T* is the period of the oscillations and 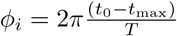 parametrizes the initial phase shift between the beginning of the transient input and the instant of the maximum in the activity just preceding *t*_0_ (this quantity is similar to the classical *phase response curve* in weakly perturbed oscillators). The asymptotic phase shift Φ_*t*_ is shown in Fig. 8C as a function of Δ*I*_*E*_ and *ϕ*_*i*_. As expected, for small transient pulse amplitude, a small phase shift arises regardless of the timing. However, when Δ*I*_*E*_ increases, an unstable region appears, and the phase response becomes much more complex and varies sharply as a function of both parameters (details of the dependence are not considered here, the point being that a strong dependence arises before the switch).

**Figure 8:**
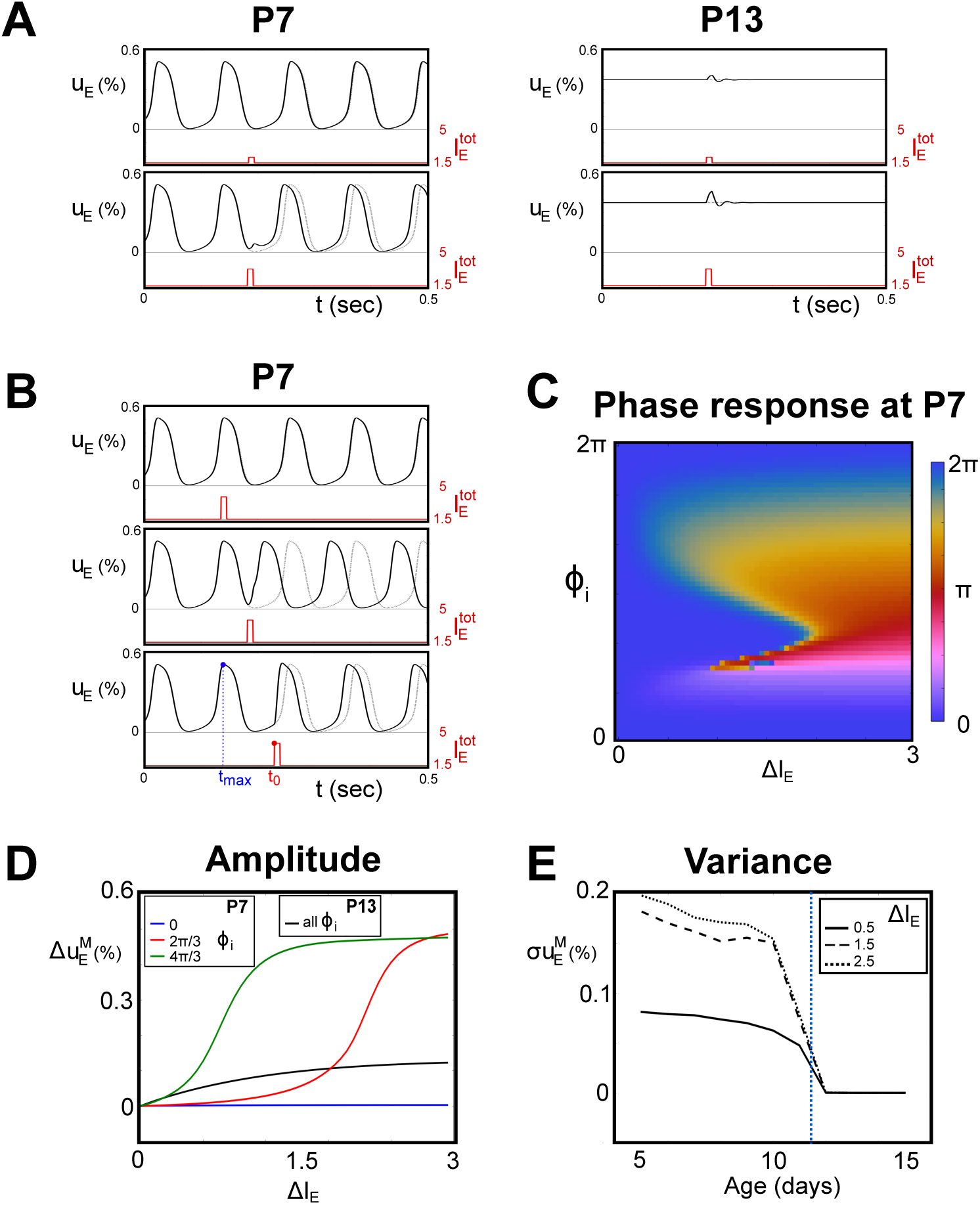
Response to external input transients. We consider the response of the excitatory population when a 10ms square input is applied to both excitatory and inhibitory populations. Panel A: results corresponding to P7 (left) and P13 (right), where model parameters are fixed as in Fig. 2 and Fig. 6, in absence of noise, for two different external stimuli, Δ*I*_*E*_ = 0.5 (top) and Δ*I*_*E*_ = 1.5 (bottom). Solid black lines represent *u*_*E*_ when an external input transient is applied (red solid lines), while dotted ones are the corresponding activity 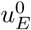 for baseline constant input. Panel B: responses at P7 for different onset time *t*_0_ of the transient external stimuli, relatively to the instant of previous maximum of activity *t*_max_. Panel C: the colormap represents the asymptotical phase shift Φ_*t*_ between the excitatory activity response and the unperturbed solution, as a function of the transient amplitude Δ*I*_*E*_ and the initial phase shift *ϕ*_*i*_. Panel D: maximum difference in activity between perturbed and unperturbed dynamics, for different initial phase shift *ϕ*_*i*_, in a window of 100ms from the onset time of the external impulse (P7 and P13 model parameters). Panel E: variance of the distribution over *ϕ*_*i*_ as a function of the age of the animal (parameters fixed as in Fig. 6), for different values of Δ*I*_*E*_. The blue dashed vertical line indicates the moment of the developmental switch, happening between P11 and P12.

**Figure 9:**
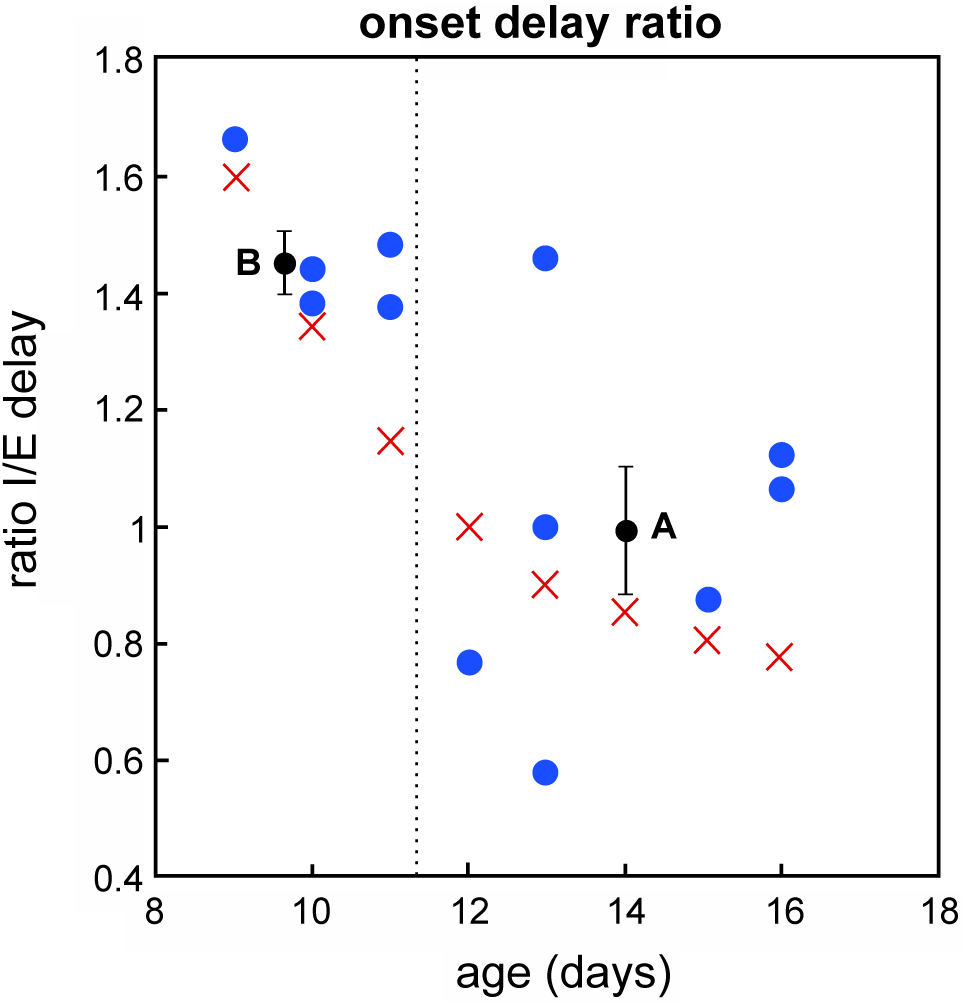
Decreasing ratio of delays. Blue points represent the experimental data of Fig. 1(C) expressed in terms of ratio between inhibitory and excitatory currents onset delays. The dashed line indicates the moment of the observed developmental switch, between P11 and P12. Black points and error bars represent the mean values and standard error for the points before (B) and after (A) the transition. Red crosses corresponds to the trajectory for *κ* chosen in the example of Fig. 6.

This dependence in the timing of the stimulus presentation is further apparent in Fig. 8D-E, when considering the maximal difference between perturbed and unperturbed activity in a window Δ*t* = 100*ms* induced by impulse stimuli at different phases (Fig. 8D) before the switch (P7):

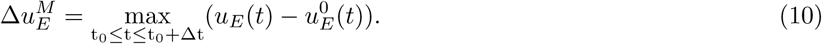

Depending on *ϕ*_*i*_, very different non-linear responses can be recorded. Because of the absence of spontaneous oscillations, no such variability arises after the switch. The switch is thus associated with a dramatic drop in the variance 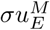 of the response distribution as a function of the time of presentation and at all stimulus amplitudes as we cross the developmental transition (Fig. 8E). After the developmental switch (P≥12), only a transient response over the asymptotic stable solution appears, and its amplitude gradually increases with that of the external transient stimulus, independently on the precise parameters corresponding to different days in our chosen trajectory example (Fig. 8D-E). Therefore these results show that our simple development model in terms of a decrease of the inhibitory/excitatory onset delay *κ* is compatible with conditions (c5) and (c6).

## Discussion

During thalamocortical development, multiple aspects of network activity undergo dramatic changes at all scales, from microscopic to macroscopic, during spontaneous activity as well as in response to external stimuli.

Using nonlinear dynamical systems theory we studied how underlying synaptic and circuit changes, generally slow and progressive, can give rise to abrupt transitions in the global network dynamics. In particular, we focused on one specific phenomenon, the sharp switch occurring in the rodent visual cortex, which changes the pattern of spontaneous activity in under 12 hours. This switch transitions a regime in which 5-20 Hz oscillations, called ‘spindle-bursts’ or ‘delta-brushes’, are produced in response to thalamocortical input, to a regime in which such input generates asynchronous irregular firing.

In order to infer the possible mechanism inducing such a transition, we developed a new version of the standard Wilson-Cowan model for excitatory and inhibitory population activities, enriched with double-exponential synapses. This modification allowed us to introduce new features relevant to developing neurons, specifically modulating the delay of each population’s response to input. These timescales indeed appear particularly prominent, as inhibitory and excitatory cells display a significant maturation of the profile of the synaptic responses to current pulses throughout the switch (Colonnese and Phillips 2018).

The Wilson-Cowan approach has been extensively studied in the case of mature cortical networks (Ermentrout 1998; Destexhe and Sejnowski 2009), and one of its major advantages is its simplicity, yet ability to accurately reproduce observed behaviors. Indeed, only a few biologically-related parameters define the model, a massive simplification compared with more biologically accurate models. If, on one hand, this obscures a direct relationship with experimental measurements, it provides a direct understanding of the main rate-based mechanisms underlying the global dynamics of the system. In (Rahmati et al. 2017) for example, another version of the Wilson-Cowan model, enhanced by short-term plasticity, has been used to address another related aspect of the development, namely the emergence of sparse coding. In this work, we have shown that the decay of the inhibitory onset timescale, which we examined as ratio to the excitatory timescale, explains the main features of the developmental switch observed experimentally. Mathematically, when the ratio between the two onset delays decreases, the dynamics of the system crosses a Hopf bifurcation arising where excitatory and inhibitory delays are of the same order of magnitude. Moreover, the model constrains directly the possible changes in electrophysiological parameters consistent with the experimental observations. In particular, we showed that an acceleration of the early oscillations and a decay of their amplitude, which occurs experimentally before the switch, predicts specific changes in timescale ratios and amplitudes. Namely, it predicts a decay of the onset delay ratio of inhibitory to excitatory currents and an increase in the relative inhibitory current amplitude. Both of these key phenomena were observed experimentally during development (Fig. 1).

To exhibit this result, we provide an extensive numerical analysis of the codimension-one and -two bifurcations occurring as a function of the normalized inhibitory delay together with relative inhibitory strength and the amplitude of the external input. We found that smooth progression in these parameters caused a flip from an initial oscillatory state to a constant activity regime. Generally, this switch could be driven either by increasing the inhibitory amplitude or by reducing the inhibitory response time. However, the experimentally measured acceleration of frequency and reduction in amplitude can only be accounted for by significant changes in the inhibitory delay, suggesting it is the key parameter driving development.

Because of the natural biological variability of the parameters, it is a complex task to assess whether changes in the electrophysiological parameters are sudden or continuous. We argue that our mathematical model and analysis is relevant in both cases: in a system that does not display the bifurcation related to timescale ratios, even a sharp decrease in the inhibitory delay would not result in the observed qualitative change in network behavior because of the persistence of a stable attractor dynamic of the system.

From the biological viewpoint, this result suggests that the transition is intrinsically related to the fact that the inhibitory population response time become faster by a larger fraction than the excitatory response time. In other words, the inhibition speeds up fractionaly faster through the switch than the excitation. This is easy to see; if we denote 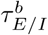 and 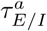 as the onset timescales of the E/I populations before and after the switch respectively, a decreasing ratio inhibitory/excitatory (*κ*) gives

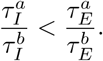

Notice that this is result is independent of the fact that 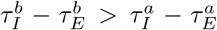 already observed in Fig. 1(C) and which could theoretically occur with constant, increasing or non-monotonic evolutions of the ratios. Going back to experimental data, we computed the ratio of the excitatory and inhibitory onset delay, and found preliminary evidence of the decay of ratios according to the above formula (Fig.9). The data indeed shows a significant decrease of the timescale ratio before the switch *κ*^*b*^ = 1.47 *±* 0.05, and after the switch of *κ*^*a*^ = 0.98 *±* 0.11 (mean *±* standard error, one-tail Welch’s t-test for *κ*^*b*^ > *κ*^*a*^ yields a p-value: *p* = 0.001). However, due to the limited number of points, more data need to be collected specifically for this purpose in order to resolve the fine evolution of the ratio as a function of age.

While not designed to model the switch from discontinuous to continuous activity that occurs alongside the switch from the oscillatory to stable regimes, our results suggest that a single mechanism can cause both. Previously it has been proposed these two changes are the result of separate (though perhaps linked) circuit changes: increased inhibition drives the stabilization of thalamocortical networks and increased input to thalamus from ascending neuromodulators and/or continuous retinal input drives continuity (Colonnese et al. 2010; Colonnese 2014; Murata and Colonnese 2018). However our results show that even in the absence of changes in strength of input (imagined as *I*_*E*_ − *I*_*I*_), as long as there is some minimal excitation occurring during low activity periods we would expect a switch from persistent down-state to a bistable up/down regime at close to the same values for *κ* that we see the switch from oscillations to stable up-states. Thus periods of low input that previously resulted only in network silence now produce alternating activity and down-states, and inputs which produced oscillations now produce stable activation, which is exactly what is observed in vivo (Shen and Colonnese 2016).

Initially we aimed to account for a qualitative change in the dynamics of cortical networks when inhibition becomes faster. Yet, we were able to capture not only the main network activity features observed experimentally, but also the order of magnitude of the frequency and amplitude trajectories of the oscillations without need for fine-tuning. The exact quantitative results however depend on the actual parameter values. In particular, while the existence of the pre-swtich oscillations depends only on the ratio between the inhibitory and excitatory onset delays, the frequency of those oscillations is proportional to the absolute value of those delays. In our final quantitative simulations, we focused on the case where the excitatory current delay is kept fixed; studies based on further experimental measurements could allow consideration in more detail the role of the absolute excitatory delay and its developmental dynamics.

Our simple model can be improved in several directions. First our model does not distinguish distinct neural populations and the variety of their timescales within the thalamocortical loop. A direct perspective would be to make the model more precise by including various populations and changes in their timescales and relative impact. In particular, the early thalamocortical oscillations described in Fig. 1 involve the thalamus, itself showing an oscillatory activity at the same frequency as cortex, and experimental evidence shows that the intact cortico-thalamic loop is a critical component of these oscillations (Murata and Colonnese 2016). The model of early oscillation generation in the visual system we developed here suggests that unpatterned input (roughly *I*_*E*_ − *I*_*I*_ in the model) from retina provides the drive, and thalamus and cortex as a whole oscillate in response. This reproduces accurately the fact that both thalamus and cortex switch to asynchronous/tonic patterns of firing after the switch, and both are the sites of reduced delay and increased amplitude of inhibition (Murata and Colonnese 2018). The value of the present results lies in showing that recurrent networks of excitatory and inhibitory neurons can be made to undergo developmental transitions similar to those observed in vivo, in response to similar changes in the electrophysiological parameters. Future studies will reveal the circuit specifics of these networks during development.

A natural question arising from this study is: are there known developments of the inhibitory circuitry that could underlie the decrease in population timescale ratios predicted by our model? Evidence shows that ‘fast-spiking’ parvalbumin-expressing interneurons, which provide rapid, persistent inhibition in the cortical network (Hu et al. 2014) gradually acquire their characteristics during the second and third postnatal week (Luhmann and Prince 1991; Huang et al. 2007). Furthermore, in somatosensory cortex, inhibitory circuits undergo a dramatic rearrangement, with thalamocortical axons changing their target from slower somatostatin interneurons neurons in layer 5 to parvalbumin expressing layer 4 interneurons (Tuncdemir et al. 2016; Marques-Smith et al. 2016; Daw et al. 2007). This development of feedforward, hence faster, inhibition in cortex is accompanied by the incorporation of the inhibitory thalamic reticular nucleus within the thalamocortical loop (Murata and Colonnese 2016), which would be expected to further speed and increase the bulk inhibitory population response. Future iterations of the model could incorporate the local connectivity of each of these inhibitory subpopulations in the model, but for the current iteration, with a single rate standing in for all inhibitory types, such sequential integration of feedforward interneuron type into the network should result in an apparent quickening of the mean response as modeled here.

We should also emphasize that our extension of the Wilson-Cowan model to investigate temporal dynamics of inhibition has implications and applications beyond development. Disruption of inhibition, particularly mediated by fast-spiking parvalbumin neurons, has been implicated in an number of neuro-developmental disorders (Gogolla et al. 2009; Contractor et al. 2015). These include multiple mouse models of autism and schizophrenia. While to our knowledge, direct measurement of inhibitory delay have not been published, reduced drive (Gibson et al. 2008) or density of fast-spiking parvalbumin neurons should result in net slowing of inhibitory rise times as represented within our model. The bifurcations of this model show that even small changes in delay can result in significant changes in the oscillatory dynamics, which might inform processing deficits observed in these conditions.

## Supporting Information

### S1 Text Mathematical details and extra parameters investigation

**S1 Fig. Dependence of the response activity on parameters *κ* and *λ*’s**. (A) The response of the inhibitory activity *u*_*I*_ to a Dirac input at dimensionless time 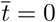, for different values of *κ*, with *λ*_*I*_ = 0.8 and *α* = 1. (B) The same response of the excitatory activity *u*_*E*_, for different values of *λ*_*E*_.

**S2 Fig. Bifurcation diagram in the space of external input** *I*_*E*_ **and currents area ratio** *α* **for different onset delays** *κ* **and different values of the connectivity parameters** *J*_*EI*_ **and** *J*_*IE*_. The limit point lines are the same for all values of *κ*. However the stability of the same limit points changes with the different values of *κ*. Solid and dotted green lines represents respectively stable or unstable Hopf bifurcations. Codimension 2 bifurcations follow the same conventions as in Fig. 4, with **ZH**: Zero-Hopf. For the clarity of the figure, saddle-node homoclinic, saddle homoclinic bifurcations and limit points of cycles are not represented here.

**S3 Fig. Dependence on the parameter** *r*. Bifurcation diagrams in 2-dimensional parameter space (*r, κ*) for fixed *I*_*E*_ = 1.5 and *α* = 1. All other parameters are fixed as specified in the main text. Color-code for areas and codimension 1 and 2 bifurcations as in Fig. 3 and Fig. 4

**S4 Fig. Dependence on the parameters** *λ*_*E*_ **and** *λ*_*I*_ : Bifurcation diagrams in 1-dimensional parameter *κ*, for fixed *I*_*E*_ = 1.5 and *α* = 1. On the left panel we compare the two symmetric cases A1: *λ*_*E*_ = *λ*_*I*_ = 0.99 and A2: *λ*_*E*_ = *λ*_*I*_ = 0.3. In the right panel, the two asymmetric cases B1: (*λ*_*E*_ = 0.3, *λ*_*I*_ = 0.9) and B2: (*λ*_*E*_ = 0.9, *λ*_*I*_ = 0.3). All other parameters are fixed as specified in the main text. Color code for areas and codimension 1 and 2 bifurcations as in Fig. 3 (L4), with light colors for cases A1 and B1, and darker colors for cases A2 and B2.

## Acknowledgments

The authors thank M. Krupa for discussions. MTC is supported by The National Eye Institute (EY022730). A.R., J.T. and B.G. acknowledge funding from ANR-10-IDEX-0001-02 PSL*. BSG was partially funded by ANR-10-LABX-0087 IEC. This study was performed as a part of the Basic Research Program at the National Research University Higher School of Economics (HSE) supported by the Russian Academic Excellence Project ‘5-100’.

## Footnotes

1 This model, while reproducing accurately the time profile of synaptic responses, does not, strictly speaking constrain the proportions to remain between 0 and 1; to overcome this issue, for each solution, we confirmed that all solutions we computed never left the interval [0, 1].

2 In fact, it can be shown (see SI text) that the peaks of the impulse responses of the excitatory and inhibitory synapses (the *onset times*) can be expressed as: 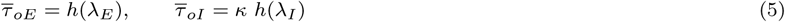 with 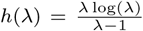. Equivalently, by reintroducing the standard time units, one has 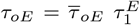 and 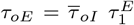, and, when *λ*_*E*_ = *λ*_*I*_, we have 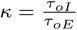.

