## Supplementary Information for "Progressive Alignment of Inhibitory and Excitatory Delay May Drive a Rapid Developmental Switch in Cortical Network Dynamics"

### Development of Inhibitory Synaptic Delay Drives Maturation of Thalamocortical Network Dynamics (Supporting Information)

Alberto Romagnoni Matthew T. Colonnese Jonathan D. Touboul Boris Gutkin

#### I Mathematical details

In this section we discuss some mathematical details concerning the theoretical model described in the main text.

Starting from the model defined in Eq. 2 in the main text, and applying the time redefinition $\bar{t} = \frac{t}{\tau_1^E}$, and the reparametrization of Eq. 4 described in the main text, we obtain the system:

$$u_E'' + \frac{1+\lambda_E}{\lambda_E} u_E' + \frac{u_E}{\lambda_E} = f_E \; (1-u_E) \; S^I(J_{EE} \; u_E + J_{IE} \; u_I + I_E),$$
$$u_I'' + \frac{1+\lambda_I}{\kappa\lambda_I} u_I' + \frac{u_I}{\lambda_I \kappa^2} = f_I \; (1-u_I) \; S^I(J_{EI} \; u_E + J_{II} \; u_I + rI_E). \quad \text{(S1)}$$

In order to understand the role of the parameters $\kappa, \alpha$ and $\lambda$'s discussed in this work, we consider here the case in which the r.h.s. of the differential equations are substituted by Dirac external impulses at time $t_0$:

$$(1-u_E(\bar{t})) \; S^I(J_{EE} \; u_E(\bar{t}) + J_{IE} \; u_I(\bar{t}) + I_E) \to \delta(\bar{t} - \bar{t}_0),$$
$$(1-u_I(\bar{t})) \; S^I(J_{EI} \; u_E(\bar{t}) + J_{II} \; u_I(\bar{t}) + rI_E) \to \delta(\bar{t} - \bar{t}_0). \quad \text{(S2)}$$

Therefore the simplified differential equations for the excitatory and inhibitory activities can be written as:

$$u_E'' + \frac{1+\lambda_E}{\lambda_E} u_E' + \frac{u_E}{\lambda_E} = f_E \; \delta(t-t_0),$$
$$u_I'' + \frac{1+\lambda_I}{\kappa\lambda_I} u_I' + \frac{u_I}{\lambda_I \kappa^2} = f_I \; \delta(t-t_0). \quad \text{(S3)}$$

This dynamical system can be analytically solved, and the general solutions read:

$$u_E(t) = \frac{\lambda_E \; e^{-\frac{t-t_0}{\lambda_E}} \; (c_1 + c_2 + f_E \; \theta(t-t_0)) - e^{-(t-t_0)} \; (c_2\lambda_E + c_1 + f_E\lambda_E \; \theta(t-t_0))}{\lambda_E - 1},$$
$$u_I(t) = \frac{e^{-\frac{(\lambda+1)(t-t_0)}{\kappa\lambda_I}}}{\lambda_I - 1} \Big( \lambda e^{\frac{t-t_0}{\kappa}} \; (c_2\kappa + c_1 + f_I\kappa \; \theta(t-t_0)) - e^{\frac{t-t_0}{\kappa\lambda_I}} \; (c_2\kappa\lambda_I + c_1 + f_I\kappa\lambda_I \; \theta(t-t_0)) \Big). \quad \text{(S4)}$$

where $\theta(x)$ is the Heaviside theta function, and $c_1, c_2, c_3, c_4$ integration constants. When imposing the solutions to be trivial before the external impulses $(u_E(t) = u_I(t) = 0, t < t_0)$ , we finally obtain:

$$u_E(t) = \frac{f_E\lambda_E \; \theta(t-t_0) \left( e^{-(t-t_0)} - e^{-\frac{t-t0}{\lambda_E}} \right)}{1-\lambda_E},$$
$$u_I(t) = \frac{f_I\kappa\lambda_I \; \theta(t-t_0) \; \left( e^{-\frac{(t-t_0)}{\kappa}} - e^{-\frac{t-t_0}{\kappa\lambda_I}} \right)}{(1-\lambda_I)}. \quad \text{(S5)}$$

It is easy to verify that the area under the curves of these responses are:

$$A_E = \int_{-\infty}^{+\infty} u_E(t)dt = f_E\lambda_E,$$
$$A_I = \int_{-\infty}^{+\infty} u_I(t)dt = f_I\kappa^2\lambda_I. \tag{S6}$$

In the main text we fixed the parameters $f_E = 1/\lambda_E$ in order to have an area $A_E = 1$, and we defined the ratio $\alpha = A_I/A_E = A_I$. The parameter $\alpha$ can then be identified as the quantity of response of the inhibitory population to a spike-like input, with respect to the excitatory response. Moreover, from these explicit solutions it is easy to derive the exact onset delays, defined as the time at which the responses reach their maximum

$$\overline{\tau}_{oE} = \frac{\lambda_E \log(\lambda_E)}{\lambda_E - 1}, \qquad \overline{\tau}_{oI} = \kappa\,\frac{\lambda_I \log(\lambda_I)}{\lambda_I - 1} \tag{S7}$$

In Fig. S1 we have shown the effect of changing the parameters $\kappa$ and $\lambda$'s on these solutions. These results justify our interpretation of $\kappa$ as representing the ratio between the onset delays of excitatory and inhibitory responses (which is an exact results when $\lambda_E = \lambda_I$), and as $\lambda$'s as the parameters regulating the decay slopes of these responses.

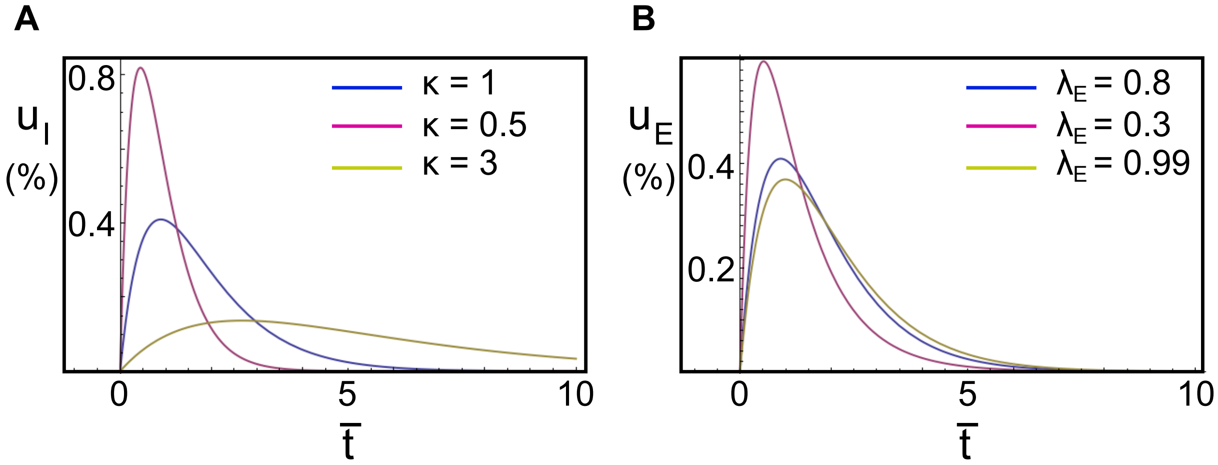


Figure S1: **Dependence of the response activity on parameters $\kappa$ and $\lambda$'s**. (A) The response of the inhibitory activity $u_I$ to a Dirac input at dimensionless time $\overline{t} = 0$, for different values of $\kappa$, with $\lambda_I = 0.8$ and $\alpha = 1$. (B) The same response of the excitatory activity $u_E$, for different values of $\lambda_E$.

#### II Independence of the results on other parameters

In the analysis discussed in the main text, we focused on the role played in our model by the three parameters $\kappa$, the ratio between the onset delays of inhibitory and excitatory currents, $\alpha$, the ratio between the inhibitory and excitatory responses, and $I_E$, the amount of external input. We fixed all other parameters of the model to standard values, and observed that a decrease in $\kappa$ is fundamental in explaining the features of the developmental switch: it determines a phase transition from an oscillatory regime to a stable one, with an increase in frequency and decrease in amplitudes, of the oscillations, when approaching the switch. In this section we show that the scenario we discussed in the main text is not peculiar of the parameters choice we made, that no particular tuning is needed, and that instead, the qualitative results are robust when the other parameters are let vary in suitable intervals.

#### II.I Connectivities

In order to show how this bifurcation diagram changes as function of the connectivity, as an example, we show in Fig. S2 the comparison between the choice we made in the main text ($J_{EI} = -J_{IE} = 10$) and the case with $J_{EI} = -J_{IE} = 9$. Limit points and Hopf bifurcations are shown for different values of $\kappa$. Notice that while LP lines are left invariant by changes in $\kappa$, Hopf bifurcations (and then the parameter region giving rise to stable oscillations) shrink when decreasing $\kappa$. Nonetheless, the dependence of the dynamics on $J_{EI}$ and $J_{IE}$ is very similar in the two cases, only the values of the parameters for which transitions happen change. For example, in the case $J_{EI} = -J_{IE} = 9$, if fixing $I_E = 1.5$, the scenario of Fig. 3(L4) in the main text cannot be reproduced for $\alpha = 1$, but instead for $\alpha \gtrsim 1.5$ for some different critical value of $\kappa$ at the Hopf bifurcation. This general behavior is guaranteed for a significant interval of values for the connectivity parameters, excluding therefore the fact that our results would depend on some sort of fine tuning. Moreover, it is easy to show that also the balanced condition $J_{EI} = -J_{IE}$ is unnecessary, and it was chosen only to diminish the number of free parameters of the model.

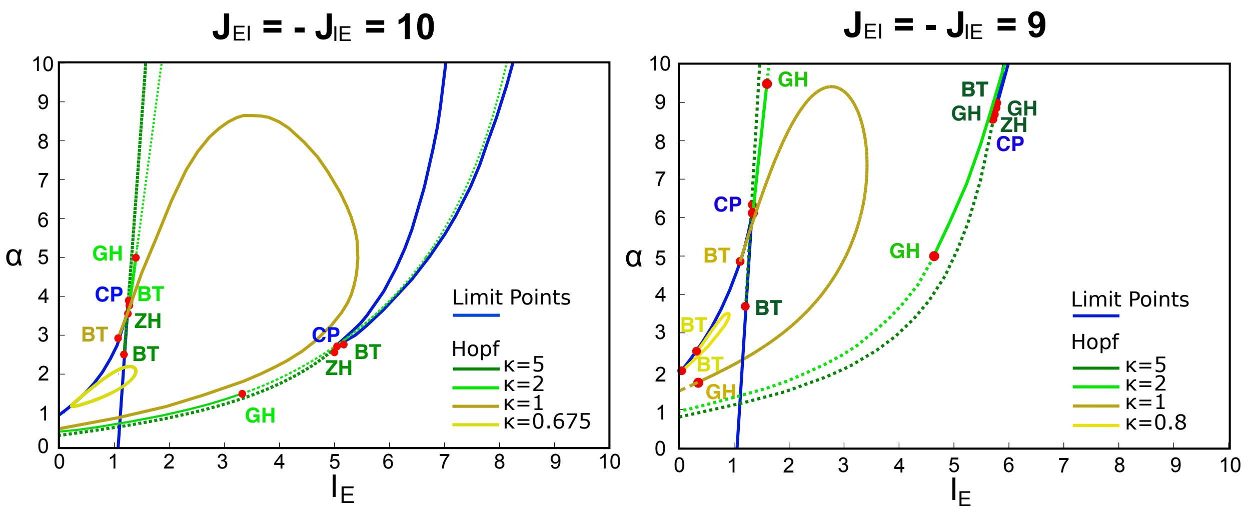


Figure S2: **Bifurcation diagram in the space of external input $I_E$ and currents area ratio $\alpha$ for different onset delays $\kappa$ and different values of the connectivity parameters $J_{EI}$ and $J_{IE}$.** The limit point lines are the same for all values of $\kappa$. However the stability of the same limit points changes with the different values of $\kappa$. Solid and dotted green lines represents respectively stable or unstable Hopf bifurcations. Codimension 2 bifurcations follow the same conventions as in Fig. 4 of the main text, with **ZH**: Zero-Hopf. For the clarity of the figure, saddle-node homoclinic, saddle homoclinic bifurcations and limit points of cycles are not represented here.

#### II.II Ratio between external inputs to inhibitory and excitatory populations

In the standard Wilson and Cowan model, the external input to inhibitory and excitatory populations are introduced as independent. In our notation, they are related by $I_I = rI_E$, therefore defining the parameter $r$ as the ratio between the two inputs. We consider here only the case of positive $r$. In Fig. S3 for $I_E = 1.5$ and $\alpha = 1$ we show a 2-dimensional bifurcation diagram in the $(r, \kappa)$ parameter pace. As for $J$'s, for a significant interval of values of this ratio ($0.3 \lesssim r \lesssim 1.1$), the developmental switch can occur for $\kappa$ decreasing with days, with the dynamics passing from an oscillatory to a stable regime, generically through an Hopf bifurcation. For $0 \lesssim r \lesssim 0.3$ and $1.1 \lesssim r \lesssim 1.3$, the transition can occur through Limit Point of Cycles bifurcations. For $r \gtrsim 1.3$ the transition, if present, occurs for unnatural higher values of $\kappa$.

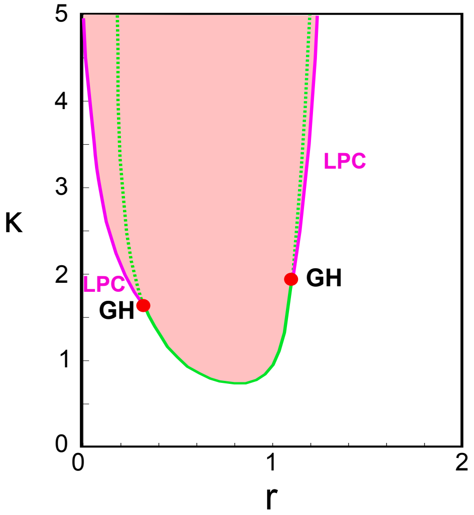


Figure S3: **Dependence on the parameter $r$.** Bifurcation diagrams in 2-dimensional parameter space ($r$, $\kappa$) for fixed $I_E = 1.5$ and $\alpha = 1$. All other parameters are fixed as specified in the main text. Color-code for areas and codimension 1 and 2 bifurcations as in Fig. 3 and Fig. 4 of the main text.

#### II.III "Decay" time constants

In the extension we propose of the standard Wilson and Cowan model with double-exponential synapses, we introduced 4 time constants (in spite of the standard 2). One was reabsorbed in the definition of time, while the others can be redefined in terms of dimensionless ratios. The main parameter $\kappa$ represents roughly the ratio between the onset delays of inhibitory and excitatory delays. The parameters $\lambda_E$ and $\lambda_I$ instead, can be seen as determining the slope of the decay of the synaptic responses and by construction they are $0 < \lambda_E, \lambda_I < 1$. In the analysis performed in the main text, we fixed $\lambda_E = \lambda_I = 0.8$. In Fig. S4 we show how once again the main qualitative result does not depend on the precise value of $\lambda$'s. In fact, in both symmetric and asymmetric cases, only the critical value of $\kappa$ at the Hopf bifurcation, and the precise value of the amplitude of the oscillations is affected by the $\lambda$'s choice.

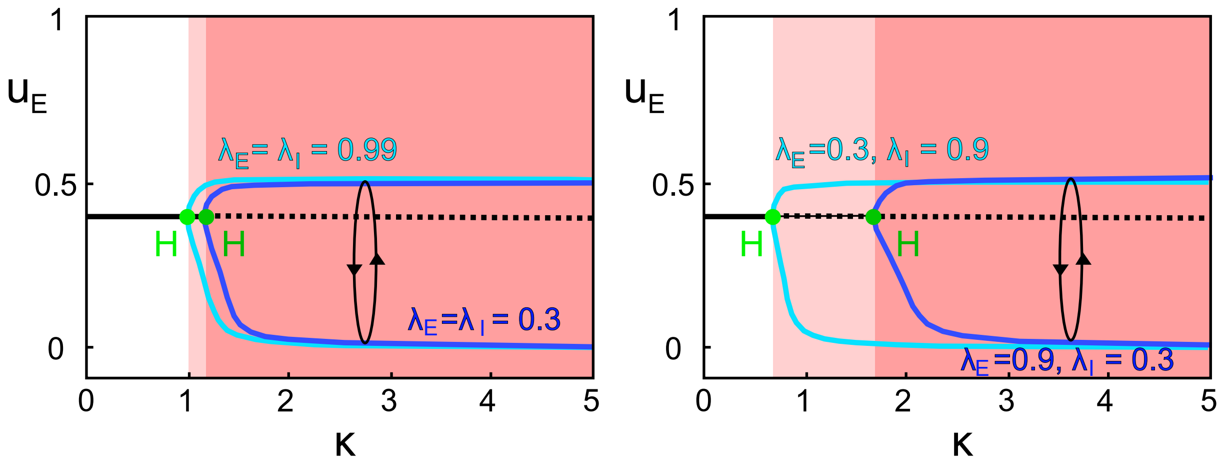


Figure S4: **Dependence on the parameters $\lambda_E$ and $\lambda_I$:** Bifurcation diagrams in 1-dimensional parameter $\kappa$, for fixed $I_E = 1.5$ and $\alpha = 1$. On the left panel we compare the two symmetric cases A1: $\lambda_E = \lambda_I = 0.99$ and A2: $\lambda_E = \lambda_I = 0.3$. In the right panel, the two asymmetric cases B1: $(\lambda_E = 0.3, \lambda_I = 0.9)$ and B2: $(\lambda_E = 0.9, \lambda_I = 0.3)$. All other parameters are fixed as specified in the main text. Color code for areas and codimension 1 and 2 bifurcations as in Fig. 3 (L4) of the main text, with light colors for cases A1 and B1, and darker colors for cases A2 and B2.
